# Highly concentrated trehalose induces transient senescence-associated secretory phenotype in fibroblasts via CDKN1A/p21

**DOI:** 10.1101/2021.10.06.463325

**Authors:** Jun Muto, Shinji Fukuda, Kenji Watanabe, Xiuju Dai, Teruko Tsuda, Takeshi Kiyoi, Hideki Mori, Ken Shiraishi, Masamoto Murakami, Shigeki Higashiyama, Yoichi Mizukami, Koji Sayama

## Abstract

Trehalose is the nonreducing disaccharide of glucose, evolutionarily conserved in invertebrates, but does not exist in vertebrates. The living skin equivalent (LSE) is an organotypic coculture containing keratinocytes cultivated on fibroblast-populated dermal substitutes. We demonstrated that human primary fibroblasts treated with highly concentrated trehalose promote significantly extensive spread of the epidermal layer of LSE without any deleterious effects. The RNA-seq analysis data and Ingenuity pathway analysis of the differentially expressed genes of trehalose-treated 2D and 3D fibroblasts at early time points revealed the involvement of the CDKN1A pathway, which is necessary for the marked upregulation of growth factors including DPT. By contrast, the mRNA-seq data of LSEs 2-weeks after air exposure indicated that gene expression profiles are similar for untreated and trehalose-treated cells in both keratinocytes and fibroblasts. The trehalose-treated fibroblasts were positive for senescence-associated β-galactosidase with the significantly downregulated expressions of LMNB1. Finally, we demonstrated that transplantation of the dermal substitute with trehalose-treated fibroblasts accelerated wound closure and increased capillary formation significantly in the experimental mouse wounds *in vivo*. These data indicate that high-concentration trehalose can induce the beneficial senescence-associated secretory phenotype in fibroblasts via CDKN1A/p21, which may be therapeutically useful for optimal wound repair.

## INTRODUCTION

Trehalose (α-D-glucopyranosyl α-D-glucopyranoside) is the nonreducing disaccharide of glucose, evolutionarily conserved in eukaryotes, plants, and invertebrates, but does not exist in vertebrates (Elbein, 1974). A pivotal method of synthesis that has dramatically reduced the production cost (Ohtake and Wang, 2011). Trehalose has been demonstrated as multifunctional and was utilized to stabilize lipids, proteins, enzymes, and tissues (Ohtake and Wang, 2011). The organ preservation solution, which was named extracellular-type trehalose-containing solution, was demonstrated to be more effective in preserving lung quality after clinical lung transplantation compared with the primary solution (Yokomise et al., 1995). Tanaka *et al*. reported that trehalose palliates the polyglutamine-mediated pathology of Huntington disease in mouse models (Tanaka et al., 2004). Trehalose inhibits the proliferation of fibroblasts owing to its inhibition of fibroblast transformation into myofibroblasts (Takeuchi et al., 2010). Additionally, a reduction in insulin/IGF-1-like signaling extends the life span of fibroblasts through an aging-suppressor function (Honda et al., 2010).

Bioengineered cellularized skin substitutes are frequently used in clinical applications as an alternative grafting technique to autografting and as *in vitro* study models, although most of the currently available models are epidermal sheets that only repair wounds by inducing keratinocytes rather than guiding a regeneration process. Multi-layered living skin equivalents (LSEs) containing bi-layered constructs that model the epidermal layer with the differentiated keratinocytes on the dermal substitutes cultivated with the fibroblasts had been used to treat skin ulcers of burn injury or epidermolysis bullosa (Kirsner, 1998). The fibroblasts in the dermal matrix of LSEs drive epidermal proliferation and differentiation through reciprocal action (Andriani et al., 2003). Numerous biomedical materials, such as type I collagen, acellular human dermis, collagen-glycosaminoglycan matrices, human plasma, and fibrin glue have been applied as dermal matrix alternatives. Nevertheless, an ideal matrix that is beneficial, readily available, and has minimal toxicity is yet to be discovered (Randall et al., 2018). The effect of trehalose on fibroblasts for LSE development remains elusive. During the course of the trials, we unexpectedly found beneficial effects of trehalose for LSE development.

Cellular senescence has been reported as a stress response in which cells experience stable cell cycle arrest following stress-inducing stimuli. The most conventional senescence marker is senescence-associated β-galactosidase (SA β-gal) activity detected after an increase in lysosomal content (Kurz et al., 2000). These senescent cells maintain metabolic capabilities and feature a hypersecretory phenotype termed the senescence-associated secretory phenotype (SASP) (Coppe et al., 2010). Although reports have characterized SASP in various cell types, its detailed composition remains unclear. The SASP is composed of a collection of proinflammatory cytokines, chemokines, and growth factors, such as epiregulin (EREG), FGF2, and VEGF (Coppe et al., 2010). The transient initiation of senescence is beneficial and contributes to the cutaneous wound repair process (Jun and Lau, 2010). Transient senescence seemed to be restricted in fibroblast-like cells, which produce platelet-derived growth factor-A (PDGFA)-enriched SASP to facilitate cutaneous wound healing (Demaria et al., 2014). In contrast, fibroblasts in which senescence is induced by oncogenic RAS oversecrete more granulocyte macrophage colony-stimulating factor (GM-CSF) and IL-6, but not EREG or vascular endothelial growth factor (VEGF), than cells in which senescence is induced by other means such as X-irradiation (Coppe et al., 2010). Basisty *et al*. reported the “SASP Atlas,” which is a comprehensive proteomic database of soluble proteins originating from multiple senescence inducers and cell types, as well as other candidate biomarkers of cellular senescence that include growth/differentiation factor 15 (GDF15), stanniocalcin 1 (STC1), and SLC1A5 (Basisty et al., 2020). Furthermore, Lamin B1(LMNB1) loss is a robust marker of senescence. There is a decline in *LMNB1* mRNA levels during senescence due to a decrease in *LMNB1* mRNA stability (Freund et al., 2012).

Cell cycle arrest is another feature of senescent cells that is controlled by activation of p53 antiproliferative function. The most pertinent function of p53 in senescence is the acceleration of cyclin-dependent kinase inhibitor 1A (*CDKN1A*) transcription (Herranz and Gil, 2018). The *CDKN1A* gene is a major target of the p53 transcription factor, and its product, p21, is a cyclin-dependent kinase inhibitor, which induces cell cycle arrest (Bartek and Lukas, 2001). In a p53- induced senescence model, Akt activation and cooperation between p21 and Akt were mandatory for cellular senescence phenotype induction (Kim et al., 2017). Polo-like kinase 1 (PLK1) is a key molecule in the G2/M transition. The induction of CDKN1A rapidly decreases cellular levels of the *PLK1* promoter activity (Zhu et al., 2002). Furthermore, high levels of p21 induce G2 arrest in normal human fibroblasts (Baus et al., 2003).

In this study, we investigated the effect of trehalose on fibroblasts mixed in type 1 collagen gel and tested whether it could affect LSE construction. We performed RNA-seq of the treated fibroblasts. Subsequently, the therapeutic potential of trehalose-treated fibroblasts in the dermal substitute as a biological dressing was investigated via skin grafting onto the full-thickness wounds of BALB/cAJc1-nu nude mice *in vivo*. These results provide an avenue for the development of a novel organotypic skin culture system for future therapeutic exploitation.

## RESULTS

### Rapid spread of LSEs containing trehalose in the fibroblast-populated collagen gel

LSEs have been used to treat skin defects. However, production of LSEs takes approximately 4 weeks, which makes LSE production impractical for applications in regenerative medicine. To investigate the beneficial effects of trehalose on fibroblasts, we constructed fibroblast-populated type I collagen gel with trehalose, upon which normal human keratinocytes were seeded to form LSEs. The sizes of the LSEs were observed after 2 weeks of airlifting at 37°C (*Figure 1A*). We evaluated the diameters of LSEs prepared in Transwell-COL with a 24-mm insert in a six-well culture plate, and the diameters of LSEs prepared with trehalose were significantly larger than those of LSEs prepared without trehalose after 2 weeks of airlifting (*Figure 1, B and C*). We confirmed this phenomenon in skin fibroblasts and keratinocytes derived from cells of three other patients 1 week after air exposure (*Figure 1-figure supplement 1, A to C*).

**Figure 1.**
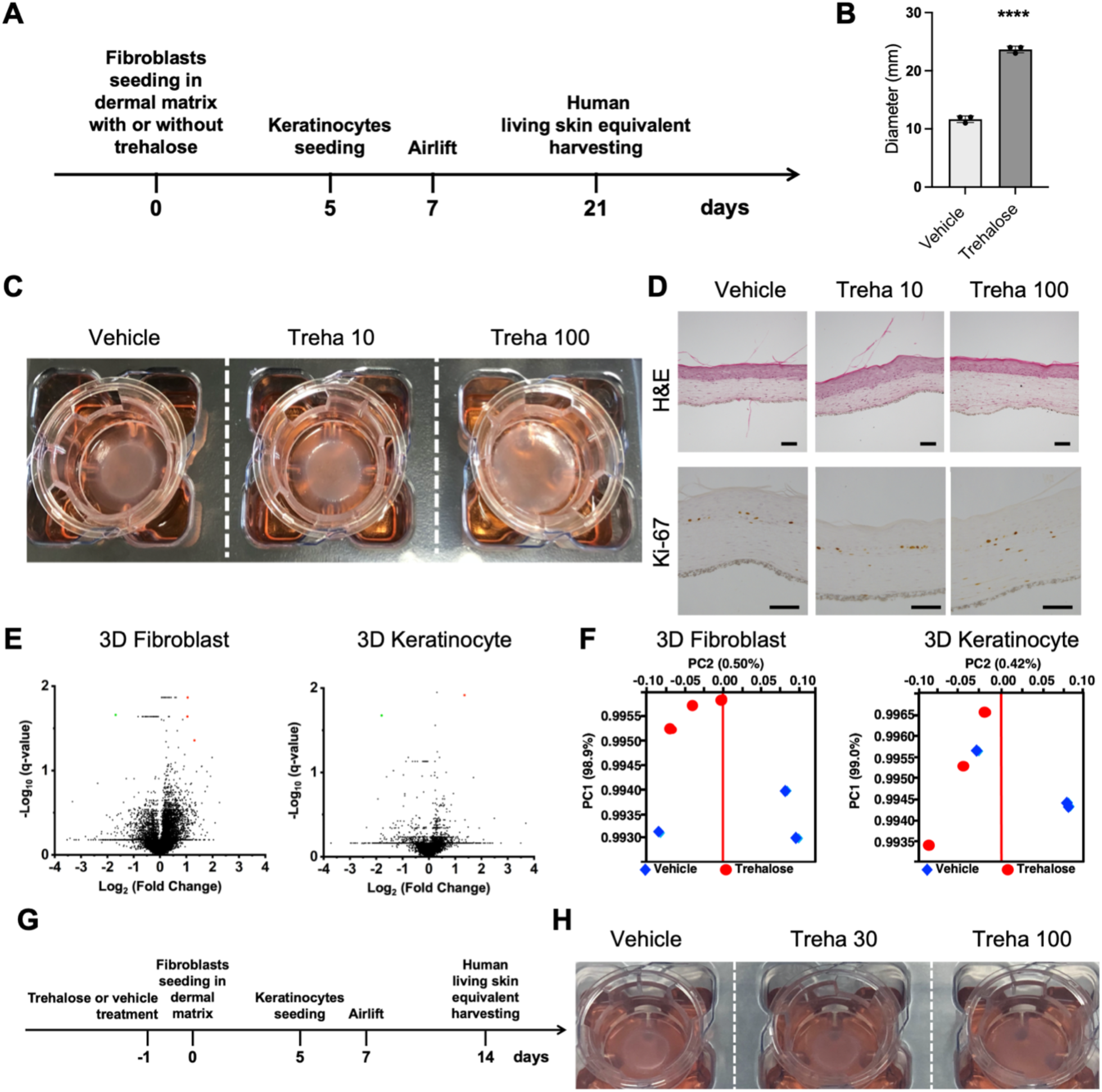
Novel effect of trehalose in the preparation of living skin equivalents. (**A**) Schematic for the preparation of cultured skin equivalents. (**B**) Diameters of LSEs with or without trehalose (100 mg/ml) added in the collagen gel, prepared in the Transwell-COL with 24- mm insert in a six-well culture plate after 2-week airlifting at 37°C. Data are expressed as means ± SD for three LSEs, which are representative of three independent experiments with similar results. ****: *P* < 0.0001 versus vehicle control groups using Student *t*-test. (**C**) Macroscopic pictures of LSEs with or without trehalose (10 and 100 mg/ml) added in the collagen gel after 2-week airlifting. (**D**) LSEs stained with hematoxylin and eosin (Scale bar = 50 μm). Paraffin-embedded sections of LSEs were sectioned and subjected to immunohistochemistry with Ki67 antibody (Scale bar = 100 μm). (**E**) Volcano plots showing gene expression in the absence or the presence of trehalose. Red or green rounds indicate genes that increased by more than 2-fold or decreased by less than the half, respectively, with less than 0.05 of q-values. (**F**) A principal component analysis (PCA) with gene expressions in the absence or the presence of trehalose showed no clear separation between principal component PC1 and PC2. (**G**) Schematic for the preparation of cultured skin equivalents with fibroblasts treated with or without trehalose before seeding in the dermal matrix. (**H**) Representative picture of LSEs with the fibroblast treated with or without trehalose (30 and 100 mg/ml) before seeding in the collagen gel after 1-week airlifting at 37°C. Data are representative of three independent experiments.

Two weeks after airlifting, hematoxylin and eosin staining was used to compare the LSEs containing trehalose (10 or 100 mg/ml) with the control LSEs. Interestingly, they were morphologically indistinguishable besides the size of the final products (*Figure 1D*). Next, paraffin-embedded sections of LSEs were subjected to immunohistochemistry with the Ki67 antibody to assess the proliferation of fibroblasts. Conversely, fibroblasts in the collagen gel with trehalose showed increased Ki67 positivity and proliferative capacity (*Figure 1D*). Additionally, we examined elastic and collagen fibers in the three-dimensional culture system with or without trehalose by Elastica van Gieson staining. Histological analysis revealed that collagen fiber (stained red) and elastic fiber (stained black) were morphologically similar among the three groups (*Figure 1-figure supplement 2A*). Hyaluronan can be detected histologically by using hyaluronan-binding protein (HABP). Interestingly, the HABP staining did not reveal differences in hyaluronan (HA) distribution between the LSEs with or without trehalose (*Figure 1-figure supplement 2B*). Alcian blue staining (pH 2.5) was used to visualize the formation of sulfated and carboxylated acid mucopolysaccharides and sialomucins in the LSEs (stained bule), with no significant changes observed between the three groups (*Figure 1-figure supplement 2C*). α-Smooth muscle actin (α-SMA) is used as a marker for myofibroblasts, which is a subset of activated fibrogenic cells. In the LSEs, α-SMA-positive cells were similarly densely lined at the dermal–epidermal junction of the three groups (*Figure 1-figure supplement 2D*). These data demonstrated that trehalose added to the collagen gel significantly accelerated proliferation of epidermal sheets, which are morphologically and histologically indistinguishable from vehicle-treated control LSEs.

To investigate the novel effect of trehalose further, we prepared larger LSEs using a larger culture insert (75-mm diameter), with proportionally more fibroblasts and keratinocytes. A rubber ring (8- mm interior diameter) was covered over the fibroblast-containing gel to stabilize it, and keratinocytes were seeded in the ring hole. The epidermal layer of LSEs containing trehalose (100 mg/ml) in the gel harvested after 2-week airlifting at 37°C spread markedly, and thus, the experimental LSEs were substantially larger than the control LSEs under the same conditions (*Figure 1-figure supplement 3*).

To examine the signaling pathways modulated by trehalose treatment in the 3D skin model, we comprehensively analyzed mRNA expressions in the epidermis and dermis of LSEs cultured in collagen gel containing trehalose (100 mg/ml) 2-weeks after air exposure. RNA-sequencing (RNA-seq) analysis revealed that genes significantly modulated by trehalose were undetected at 14 days in the keratinocytes except for the upregulated bone morphogenic protein 6 (*BMP6*) gene and the downregulated *LINC00302* gene (*Figure 1E*). In the fibroblasts, the gene expressions by trehalose resembled those in the vehicle cells, and only four genes (*SCARNA22*, *PTCHD4*, *RP11-137H2.6*, and *NPR3*) were significantly regulated by the addition of trehalose (*Figure 1E*). Principal component analysis (PCA) provided no major difference in the gene expression of the keratinocytes and fibroblasts by trehalose treatment (*Figure 1F*). In addition, the factor loadings of genes in PC1 and PC2 showed no effect on trehalose treatment. Our observations indicated that gene expression profiles in a long culture of 3D gels are similar for untreated and trehalose-treated cells in both keratinocytes and fibroblasts. Next, we examined whether fibroblast pretreated with trehalose before seeding in the collagen gel achieved increased proliferation of the epidermal layer of the 3D culture model (*Figure 1G*). Interestingly, LSEs with fibroblast pretreated with trehalose (30 or 100 mg/ml) accelerated the spread of the epidermal layer compared with the control LSEs (*Figure 1H*). These observations indicated that trehalose pretreatment on the fibroblast monolayer before seeding in the gel can induce significantly accelerated proliferation of the keratinocyte layer of LSE.

### Whole transcriptome analysis in trehalose-treated 2D and 3D fibroblasts

Gene expression profiles of both keratinocytes and fibroblasts in a long culture of 3D gels treated with trehalose were similar to those of the untreated cells. Thus, we comprehensively examined the transient gene expressions in the trehalose-pretreated fibroblasts using RNA-seq. Trehalose (100 mg/ml) treatment for 24 h induced upregulation of 1,256 genes (FC > 2.0, q < 0.01) and downregulation of 484 genes (FC < 0.5, q < 0.01) in the 2D culture compared with those of untreated cells. In the 3D culture, 267 genes or 332 genes were upregulated (FC > 2, q < 0.01) or downregulated (FC < 0.5, q < 0.01), respectively, by 72-h trehalose treatment (*Figure 2-figure supplement 1, A and D*). The gene expression profiles of the fibroblasts in 2D and 3D cultures were clearly separated by PC2 in PCA from those of untreated cells, indicating that trehalose affects cellular function through gene expressions (*Figure 2-figure supplement 1, B and E*). We plotted the factor loadings of the genes in PC1 and PC2 to observe the gene expressions involved in PC2 separation. As the positively contributing genes, growth factors, such as dermapontin (*DPT*), *EREG*, *FGF2*, and angiopoietin-2 (*ANGPT2*), were observed in both 2D and 3D cultures in addition to a cell cycle inhibitor, *CDKN1A* (*Figure 2-figure supplement 1, C and F*). The cell cycle-related genes, Aurora kinase A (*AURKA*), *PLK1*, and Myb proto-oncogene like 2 (*MYBL2*) negatively participated in the PC2 separation of 2D and 3D cultures treated with trehalose (*Figure 2-figure supplement 1, C and F*). Treatment with highly concentrated trehalose in the fibroblast cells suggests strict regulation of the cell cycle despite the release of various growth factors.

To elucidate the signaling pathways activated in highly concentrated trehalose-treated fibroblasts, we analyzed the interaction network with Ingenuity Pathway Analysis (IPA) using the information collected from databases on protein interactions. In the presence of trehalose, 131 genes were downregulated in common in 2D and 3D culture, and the expression patterns were shown on the heatmap, which included cell cycle-related genes such as Aurora kinase B (*AURKB*), *PLK1*, and Anillin actin-binding protein (*ANLN*) (*Figure 2, A and B*). The downregulated genes revealed the reduction of kinetochore metaphase signaling, G2/M DNA damage, and the cell cycle checkpoint in the canonical pathways (*Figure 2C*). With respect to upstream factors, asparaginase, a drug for acute lymphoblastic leukemia, was detected in the upstream analysis and is able to arrest the cell cycle. ZBTB17 is a transcriptional negative regulator in the cell cycle (*Figure 2D*). *CDKN1A* was also detected as an upstream factor based on the significant decrease of *PLK1*, *CDK1*, *CCNA2*, and *CDC25A* after trehalose treatment (*Figure 2E*). The inhibition of *CDK1* and *CCNA2* was suggested to partially induce the reduction of *MYBL2*. The 127 upregulated genes were observed in both 2D and 3D cultures (*Figure 2, F and G*), and contained *CDKN1A* was detected as an upstream factor of downregulated genes (*Figure 2D*). The pathway analysis using upregulated genes revealed the p53 signaling pathway involving in cell cycle arrest (*Figure 2H*). Inhibition of *AURK* and *ANLN* were detected as upstream factors in the upregulated genes and were also observed in downregulated genes in the presence of trehalose (*Figure 2, I and J*). We integrated the upregulated genes into the downregulated genes and analyzed the signaling pathways because the pathways detected by the upregulated genes were closely related to the pathways of the downregulated genes (*Figure 3A*). The graphical summary connected the network analysis to the cellular functions and showed that the senescence cells were activated by p53 and CDKN1A related to the cell cycle regulation and mitosis arrest (*Figure 3B*). In the network analysis, activation of DPT and VEGF were suggested to be induced by the Notch and Caspase (*Figure 3-figure supplement 1, A and B*). The mRNA expressions of genes involving cellular senescence, *CDKN1A*, and *LMNB1* were confirmed by quantitative PCR (qPCR) in 2D fibroblasts (*Figure 4A*). Western blot analysis revealed that trehalose treatment increased p21 expression and decreased lamin B in a dose-dependent manner (*Figure 4B*). The dose-dependently increased expression of p21 in the nuclei after trehalose treatment was confirmed using fluorescence microscopy (*Figure 4C* and *Figure 4-figure supplement 1*). Furthermore, mRNA expressions of genes involved in cellular functions such as cell cycle regulation, *AURKA*, *AURKB*, *PLK1*, and *MYBL2* were confirmed by qPCR in 2D fibroblasts (*Figure 4D*) and 3D fibroblasts (*Figure 4E*). Additionally, we also confirmed these effects of trehalose with the fibroblasts derived from three other patients. These findings suggested that trehalose possesses the ability to temporarily upregulate *CDKN1A* and downregulate *AURKA*, *AURKB*, *UBE2*, *PLK1*, *MYBL2*, and *LMNB1*, thus leading to cell cycle arrest and transient senescence of the fibroblasts.

**Figure 2.**
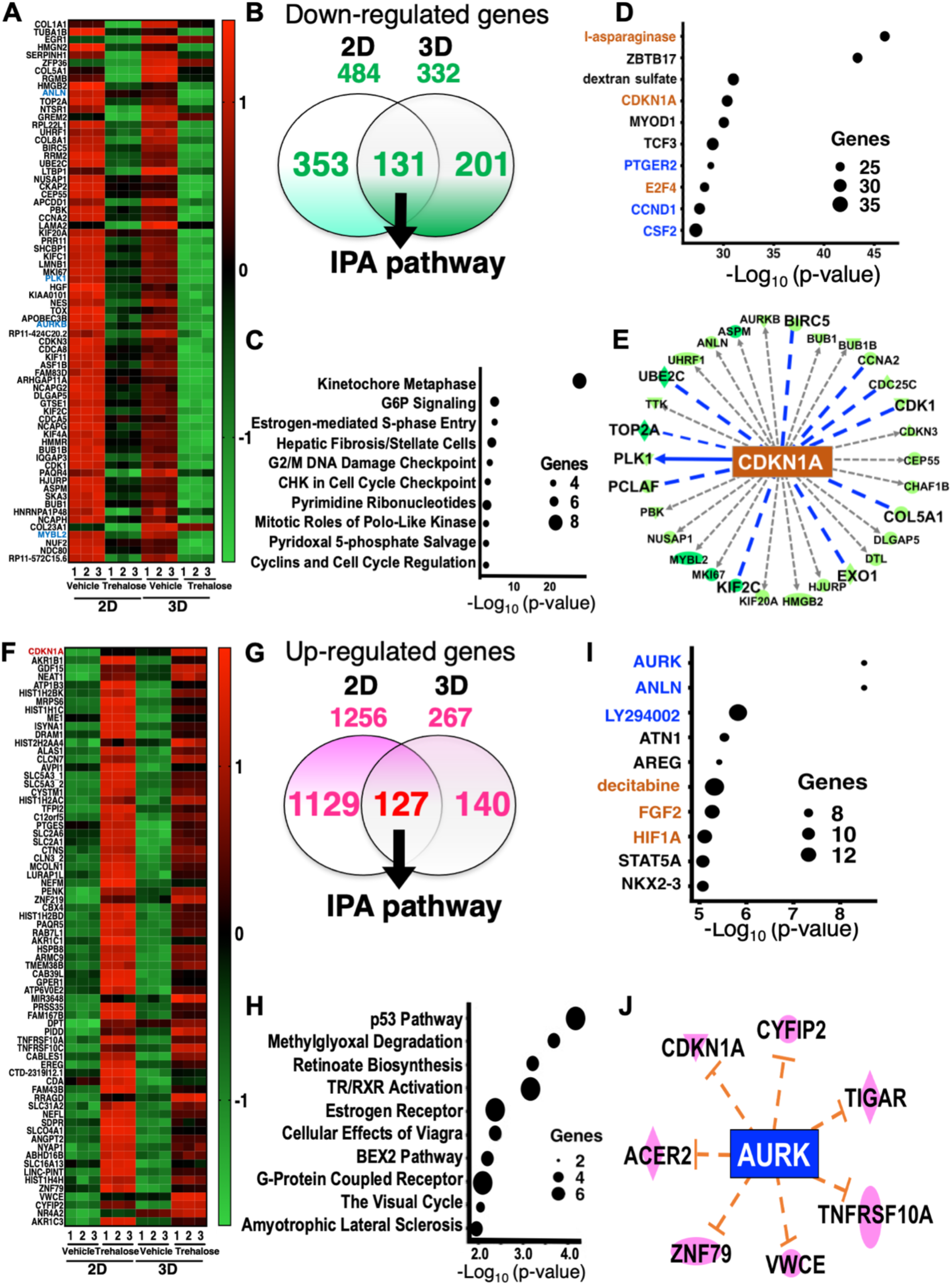
Highly concentrated trehalose-induced signaling pathways detected in human fibroblasts by whole transcriptome analysis with RNA-seq. (**A**) The heatmap shows the z-scores of the gene expression downregulated in 2D and 3D fibroblasts by trehalose. (**B**) The Venn diagram demonstrates the numbers of downregulated genes. The 131 genes downregulated in 2D and 3D fibroblasts were used for an Ingenuity pathway analysis (IPA). (**C**) Canonical pathways detected in IPA using the genes downregulated by trehalose were shown together with number of genes involved in the detected pathway. (**D**) Upstream factors of the downregulated genes by trehalose were indicated together with the number of genes involved in the detected factor. The upstream factors signaling by activation or by inhibition were shown in brown or blue, respectively. (**E**) A network demonstrates the interaction of CDKN1A, which was detected as an upstream factor of the downregulated genes, and the signals inhibited by trehalose, which are shown as blue lines. The green shapes indicate the genes downregulated by trehalose. (**F**) A heatmap shows the z-scores of the gene expressions upregulated in 2D and 3D fibroblasts by trehalose. (**G**) The Venn diagram demonstrates the numbers of the upregulated genes. (**H**) Canonical pathways detected in IPA using the genes upregulated by trehalose were shown together with the number of genes involved in the detected pathway. (**I**) Upstream factors of the upregulated genes by trehalose were indicated together with the number of genes involved in the detected factors. The upstream factors signaling by activation or by inhibition were shown in brown or blue, respectively. (**J**) A network demonstrates the interaction of AURK, detected as an upstream factor of the downregulated genes, and the factors predicted to activate interaction with AURK by trehalose, which are shown as red lines. The red shapes indicate the genes upregulated by trehalose.

**Figure 3.**
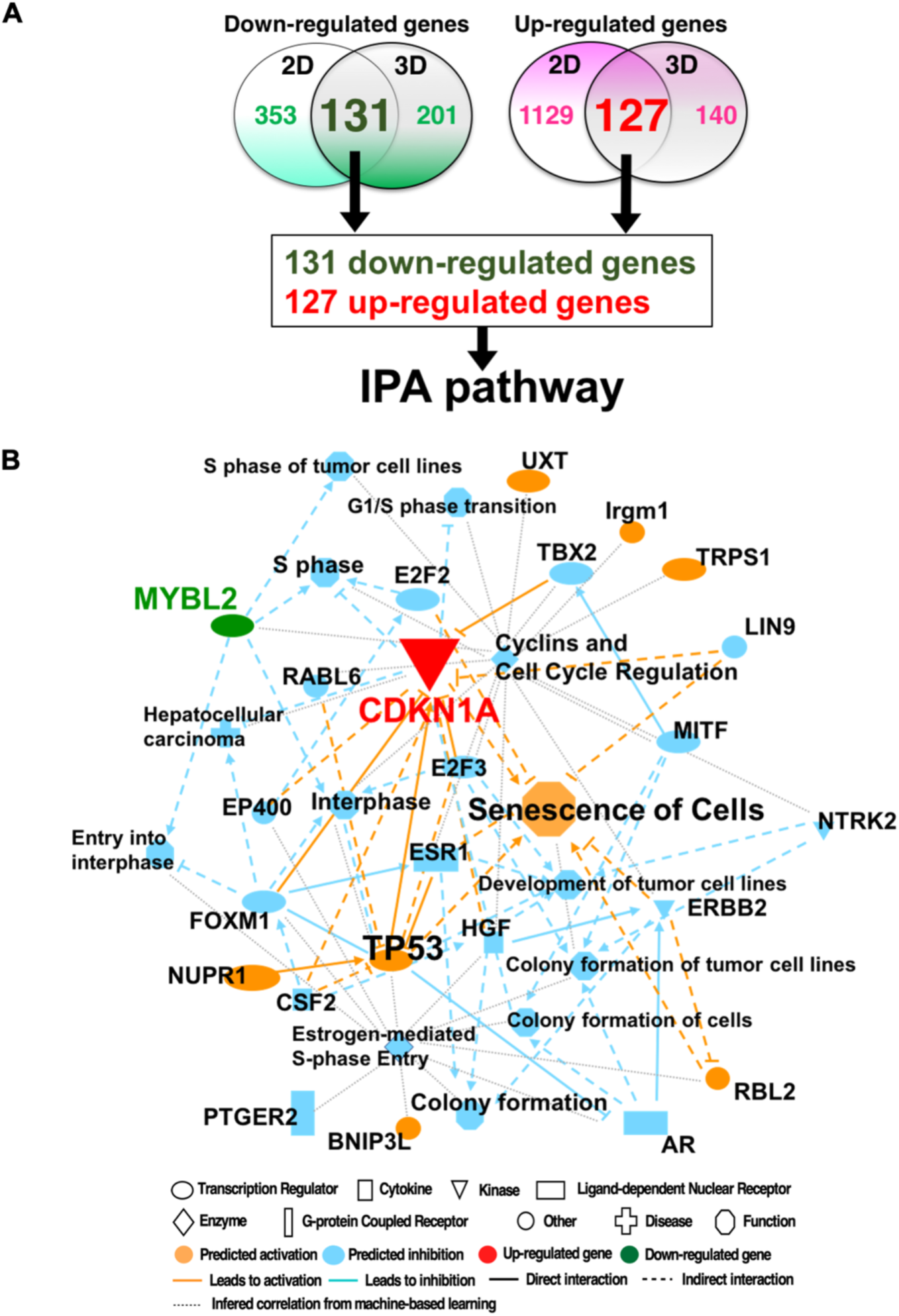
Graphical summary and network analysis by IPA pathway using genes modulated by trehalose. (**A**) Venn diagram demonstrates the numbers of the downregulated genes and the upregulated genes analyzed in Fig2. There were 131 downregulated genes and the 127 upregulated genes for an IPA. (**B**) A graphical summary shows the senescence cells, which are induced by p53 and CDKN1A, connected by the network analysis to the cellular functions.

**Figure 4.**
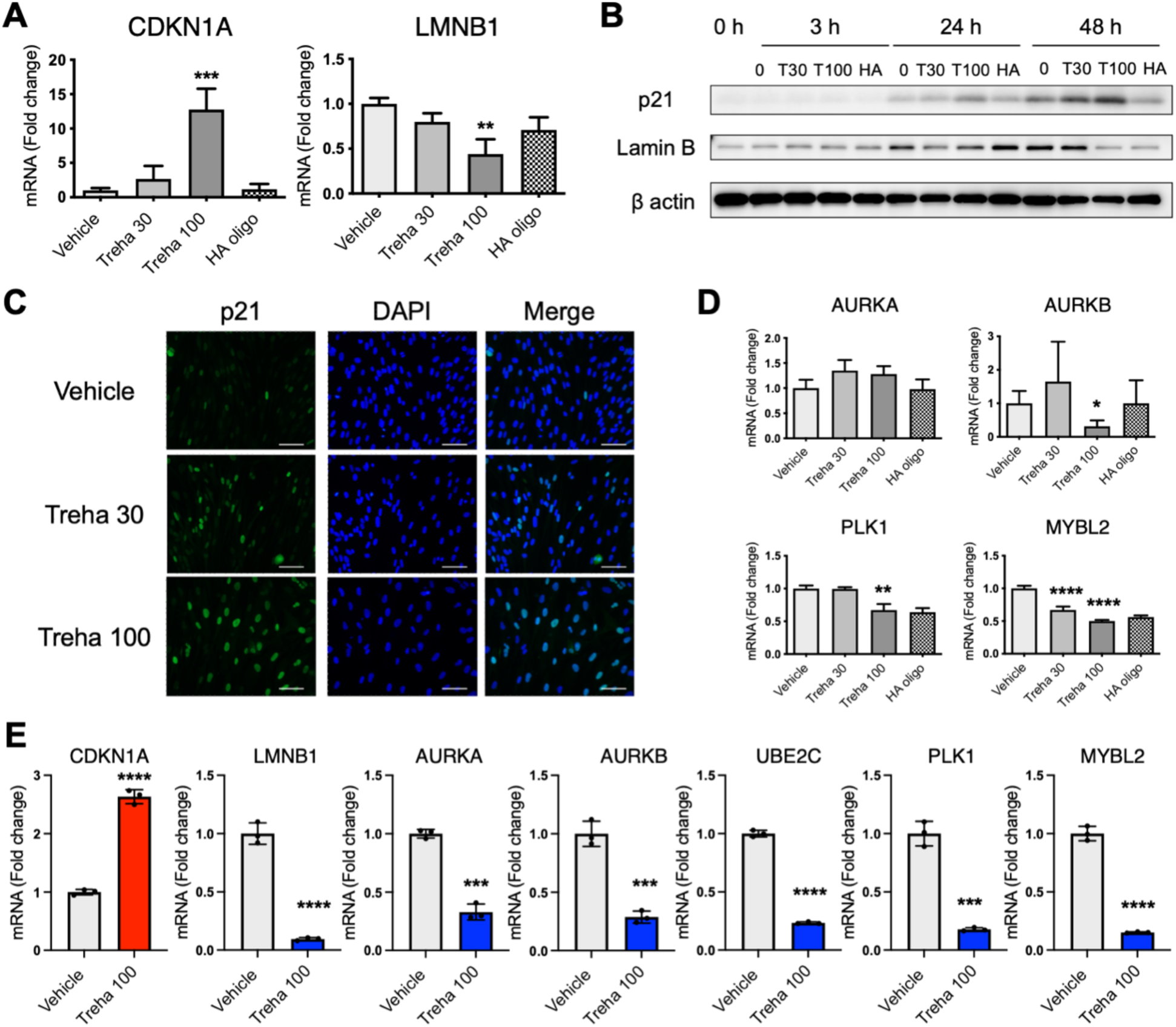
Trehalose modulates the expression of senescence and cell cycle arrest-related molecules. (**A**) Human dermal fibroblasts were treated with trehalose (30 and 100 mg/ml), tetrasaccharide hyaluronan (HA oligo) (30 μg/ml), or vehicle (PBS) for 24 h. *CDKN1A* and *LMNB1* mRNA expression were assessed by qPCR. Data are shown as relative expressions to the control (vehicle-treated) fibroblasts. (**B**) Western blotting showing the expression of p21, lamin B, and β-actin in human dermal fibroblasts treated with trehalose (30 mg/ml, 100 mg/ml) or the vehicle for 24 h. (**C**) Human dermal fibroblasts were treated with trehalose (30 and 100 mg/ml) or vehicle control (PBS) for 24 h. The cells were stained with antibody for p21 (green) and DAPI (blue) for nuclei and were observed using a fluorescence microscope. Scale bar = 100 μm. (**D**) *AURKA*, *AURKB*, *AURKC*, *MYBL2*, *PLK1*, and *UBE2C* mRNA expressions were assessed by qPCR. Data are shown as the relative expression to the control (vehicle-treated) fibroblasts. (**E**) Trehalose (100 mg/ml) or vehicle (PBS) were added in the human dermal fibroblasts populated collagen gel for 72 h. *CDKN1A*, *LMNB1*, *AURKA*, *AURKB*, *UBE2C*, *PLK1*, and *MYBL2* mRNA expressions were assessed by qPCR. Data are shown as relative expression to the control (vehicle-treated). *: *P* < 0.05, **: *P* < 0.01, ***: *P* < 0.001, ****: *P* < 0.0001 versus the vehicle-treated control group by one-way ANOVA (A, D) or Student *t*-test (E). Data are expressed as means ± SD for three wells (A, D) or three dermal substitutes (E), and are representative of three independent experiments.

### Fibroblasts in G2/M interphase and Erk1/2- and Akt-activated senescence

To further explore the effects of highly concentrated trehalose, we studied morphological alterations of fibroblasts after trehalose treatment. Phase contrast microscopy revealed dose-dependent morphological differences between fibroblasts cultured with or without trehalose. We observed that the shape of cells cultured with trehalose remained polygonal/expanded, although the control cells became fusiform/elongated (*Figure 5A and Video 1-3*). In *Figure 5A and the video 1-3*, we demonstrate that trehalose inhibited the population growth of monolayer fibroblast cells. Further, to clarify trehalose-induced cell proliferation inhibition, we examined cell viability via CCK8 assay. Trehalose slightly inhibited cellular proliferation of the fibroblasts (*Figure 5-figure supplement 1*). Importantly, instead of trehalose, we observed that a high-concentration sucrose (100 mg/ml) in the medium inhibited cell proliferation and induced cell death in human dermal fibroblasts (*Figure 5-figure supplement 1* and Video 4). Fibroblasts were characterized for a potential senescent phenotype via SA-βGAL staining, which revealed that more trehalose-treated fibroblasts than vehicle-treated fibroblasts were SA-βGAL positive (*Figure 5B*). These findings indicate that high-concentration trehalose induces the cytostatic effect and senescence.

**Figure 5.**
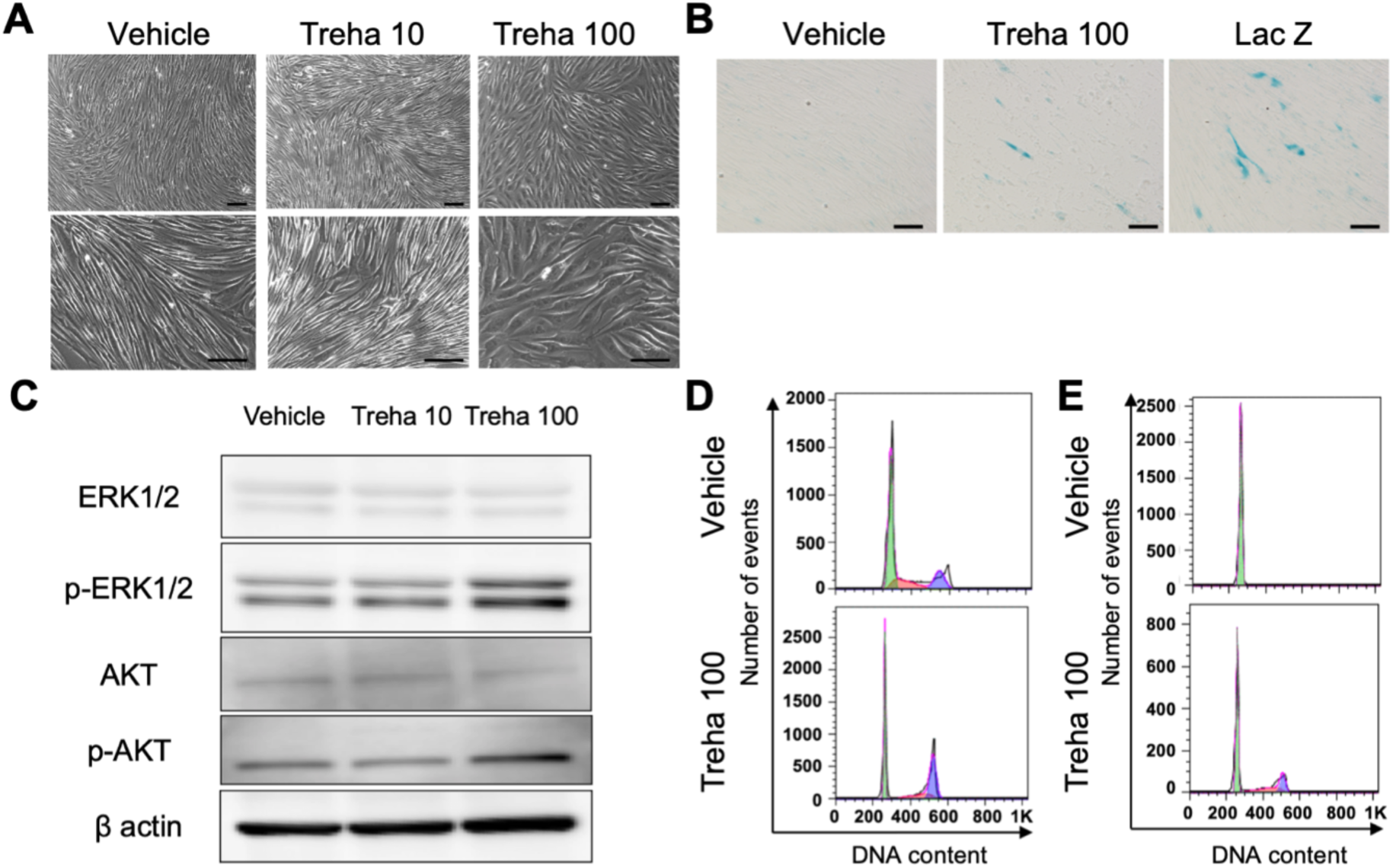
Trehalose arrests fibroblasts in the G2/M interphase of the cell cycle and triggers senescence with activation of Erk1/2 and Akt. (**A**) Representative photos of human dermal fibroblasts treated with trehalose (10 and 100 mg/ml) or vehicle (PBS) for 24 h. Phase contrast micrographs, bar = 50 μm. (**B**) Fibroblasts were characterized for a potential senescent phenotype via SA-βGAL staining after treatment with trehalose (100 mg/ml), vehicle, or Ad-LacZ (MOI:30) for 24 h. Scale bar = 50 μm. (**C**) Western blotting showing the expression of ERK1/2, p-ERK1/2, AKT, p-AKT, and β-actin in human dermal fibroblasts treated with trehalose (10 and 100 mg/ml) or vehicle for 24 h. (**D**) Twenty-four hours after the trehalose (100 mg/ml) or vehicle treatment, human dermal fibroblasts were stained with propidium iodide, and the percentage of G2/M cells was measured by flow cytometry. Representative FACS images. (**E**) Seven days after air exposure, human dermal fibroblasts in the living skin equivalent were stained with propidium iodide, and the percentage of G2/M cells was measured by flow cytometry. Representative FACS images. Data are representative of two independent experiments.

Next, fibroblasts treated with or without trehalose for 24 h were further investigated by Western blotting. High-concentration trehalose activated ERK1/2 and AKT (*Figure 5C*). To further analyze the effect of trehalose on cell cycle progression, fibroblasts treated with or without trehalose were analyzed by flow cytometry after propidium iodide staining. Of the cells treated with trehalose (100 mg/ml), 30% were arrested in the G2/M interface, whereas 20% of the cells treated without trehalose were arrested in the G2/M interface (*Figure 5D*). In addition, human dermal fibroblasts in the LSEs were stained with propidium iodide 7 days after air exposure, and the percentage of G2/M cells was measured by flow cytometry. Interestingly, 10% of cells treated with trehalose (100 mg/ml) were arrested in the G2/M interface, whereas almost none of the cells treated without trehalose accumulated in the G2/M interface (*Figure 5E*). Furthermore, there was significantly higher superoxide radical generation in those fibroblasts treated with trehalose (*Figure 5-figure supplement 2*). Therefore, we conclude that trehalose triggers two antagonistic cell cycle regulatory pathways in fibroblasts: the classical mitogenic ERK and AKT pathway and a novel G2/M cell cycle arrest pathway with induction of p21.

### Upregulation of wound healing-related genes in trehalose-treated fibroblasts

Senescent cells exhibit a hypersecretory phenotype, which has been referred to as the SASP (Coppe et al., 2010). The SASP comprises a collection of growth factors (Coppe et al., 2010, Wilkinson and Hardman, 2020). The beneficial and transient initiation of senescence could contribute significantly to a cutaneous wound healing process (Jun and Lau, 2010). Next, we aimed to elucidate the characteristics of these high-concentration trehalose-induced SASPs in human fibroblasts. RNA-seq analysis data and the IPA of the differentially expressed genes revealed significant upregulation of 10 wound healing-related genes—*EREG, ARG2, CCL2, IL1RN, PGF, SPP1, VEGFA, FGF2, ANGPT2,* and *DPT*. Then, to confirm RNA-seq findings, qPCR mRNA expression analysis of the wound healing-related genes was performed, which revealed a significant increase in fibroblasts treated with trehalose (30 or 100 mg/ml) for 24 h compared with vehicle- or HA-treated control fibroblast (*Figure 6A*). We confirmed these effects of trehalose using the fibroblasts derived from three other patients. DPT has a vital role in promoting keratinocyte migration during re-epithelialization in wound healing (Krishnaswamy and Korrapati, 2014). DPT secretion in the medium and DPT expression in the fibroblasts were assessed by ELISA and Western blotting, respectively. We found a significant increase in DPT protein secretion in the cultured medium 48 h after trehalose treatment compared with that of the vehicle-treated control fibroblasts by ELISA (*Figure 6B*). Interestingly, results from western blotting demonstrated trehalose treatment increased the remarkable DPT protein expression in cell lysate after 48 h (*Figure 6C*). Consistent with these findings, qPCR mRNA expression analysis of *DPT* in the 3D fibroblasts, which were added in the collagen gel with trehalose (100 mg/ml) for 72 h, revealed a significant increase compared with that of the vehicle-treated fibroblasts (*Figure 6D*). Furthermore, RNA-seq analysis of the 3D fibroblasts revealed 267 upregulated genes in the trehalose-treated fibroblasts compared with those of the vehicle-treated control fibroblasts, including significant upregulation of nine wound healing-related genes—*EREG, ARG2, CCL2, IL1RN, PGF, SPP1, VEGFA, FGF2,* and *ANGPT2*. Then, to confirm RNA-seq findings, qPCR mRNA expression analysis of these wound healing-related genes was performed; this revealed a significant increase in the 3D fibroblasts that were added in the collagen gel with trehalose (100 mg/ml) for 72 h compared with that of the control fibroblasts (*Figure 6D*).

**Figure 6.**
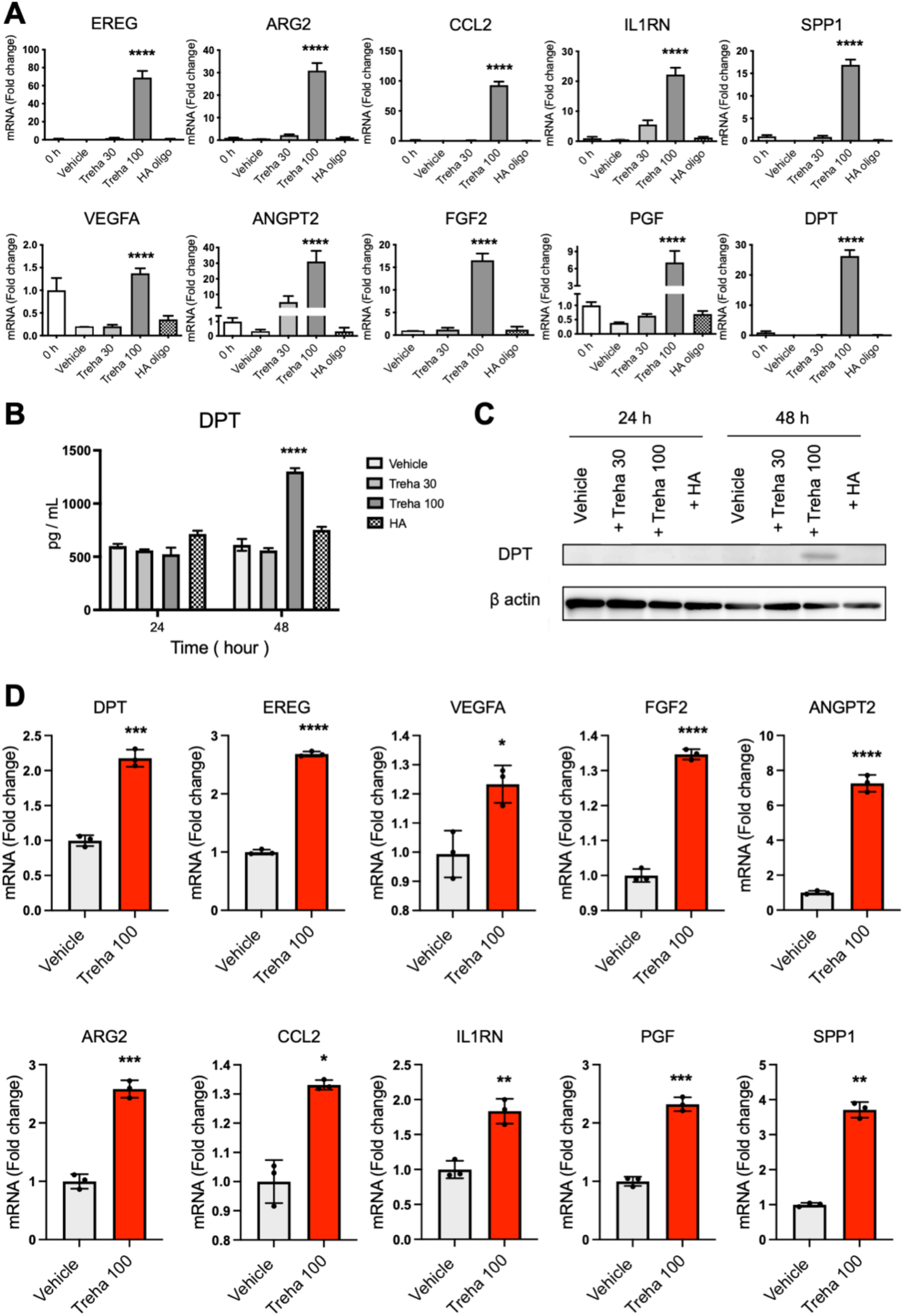
Trehalose induces an increase in the expression of wound healing-related molecules. (**A**) Human dermal fibroblasts were treated with trehalose (30 and 100 mg/ml), tetrasaccharide hyaluronan (HA oligo) (30 μg/ml), or vehicle control (PBS) for 24 h. *EREG*, *ARG2*, *CCL2*, *IL-1RN*, *PGF*, *SPP1*, *VEGF*, *ANGPT2*, and *DPT* mRNA expressions were assessed by qPCR. Data are shown as relative expression to the control (0 h) fibroblasts. For *FGF2* mRNA expression, data are shown as relative expression to vehicle control groups at 24 hours. (**B**) DPT was measured by ELISA in the culture medium of human dermal fibroblasts. One set of fibroblasts was treated with trehalose 30 or 100 mg/ml, tetrasaccharide hyaluronan (HA) (30 μg/ml), or the vehicle (PBS) for 24 or 48 h (*n* = 3). (**C**) Representative Western blots showing DPT and β-actin expression in human dermal fibroblasts 24 or 48 h after vehicle, trehalose 30 or 100 mg/ml, tetrasaccharide hyaluronan (HA) (30 μg/ml) exposure. (**D**) Trehalose (100 mg/ml) or vehicle were added in the human dermal fibroblasts populated collagen gel for 72 h. *DPT*, *EREG*, *VEGF*, *FGF2*, *ANGPT2*, *ARG2*, *CCL2*, *IL-1RN*, *PGF*, and *SPP1* mRNA expression were assessed by qPCR. Data are shown as the relative expression to the control (vehicle-treated). *: *P* < 0.05, **: *P* < 0.01, ***: *P* < 0.001, ****: *P* < 0.0001 versus the vehicle-treated control group by one-way ANOVA (A, B) or Student *t*-test (D). Data are expressed as means ± SD for three wells (A, D) or three dermal substitutes (D), and representative of three independent experiments.

### CDKN1A is involved in the upregulation of SASP factor genes

Next, we investigated the involvement of CDKN1A in trehalose-induced SASP in 2D fibroblasts. CDKN1A siRNA transfection significantly suppressed its mRNA levels, confirming successful CDKN1A knockdown (*Figure 7*). CDKN1A knockdown significantly suppressed the upregulation of *DPT*, *ANGPT2*, *VEGFA*, *EREG*, *FGF2* mRNA levels after trehalose treatment (*Figure 7*), suggesting the critical role of CDKN1A in trehalose-induced SASP in fibroblasts.

**Figure 7.**
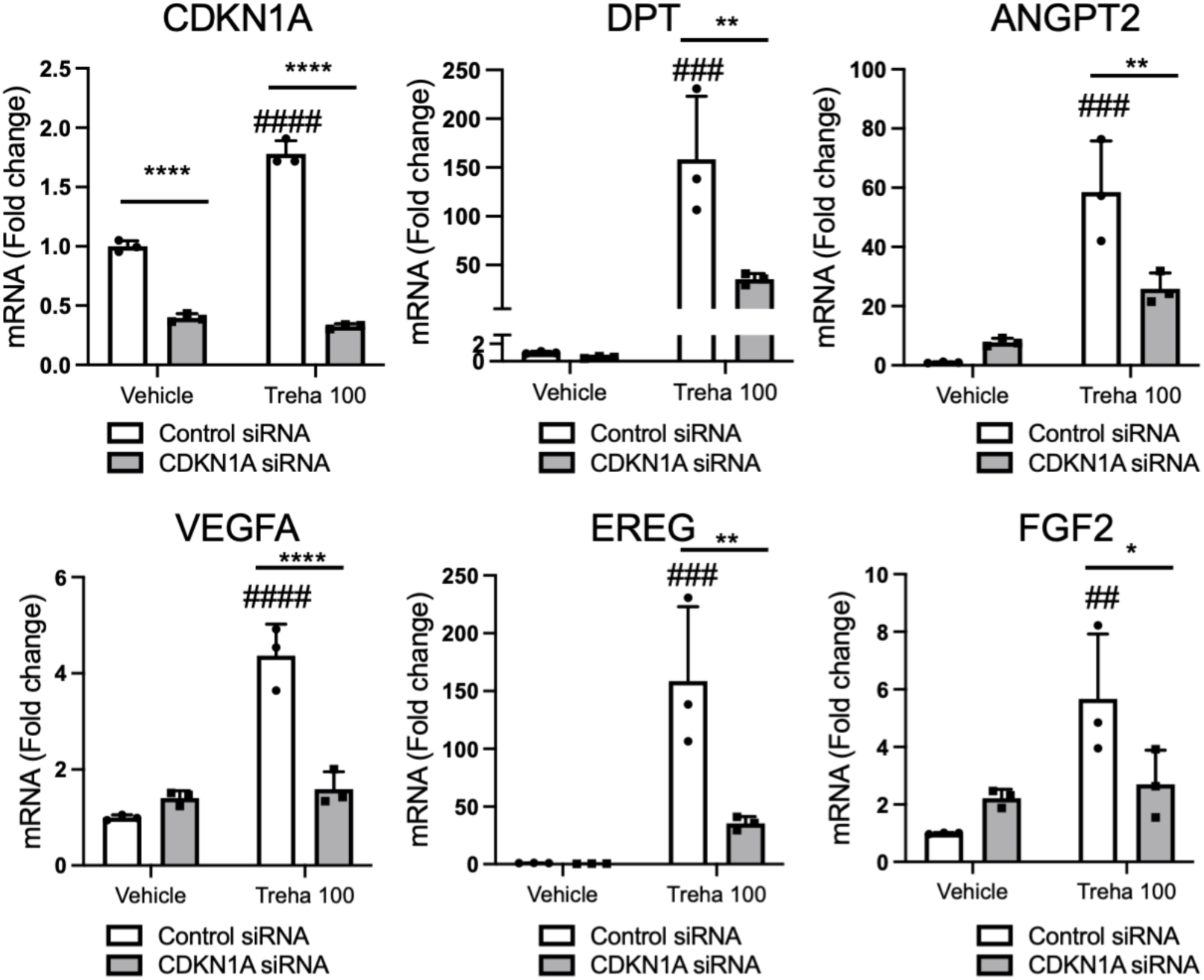
CDKN1A is involved in trehalose-induced increase in the mRNA expression of wound healing-related molecules. (**A**) After transfection with control or *CDKN1A* siRNA, human dermal fibroblasts were treated with trehalose (100 mg/ml) or vehicle control (PBS) for 48 h. *CDKN1A*, *DPT*, *ANGPT2*, *VEGF*, *EREG*, and *FGF2* mRNA expressions were assessed by qPCR. Data are shown as relative expression to the control siRNA and vehicle treated fibroblasts. Data are expressed as means ± SD, and representative of two independent experiments with similar results. *: *P* < 0.05, **: *P* < 0.01, ****: *P* < 0.0001 versus the relevant control group, and ##: *P* < 0.01, ###: *P* < 0.001, ####: *P* < 0.0001 versus the control siRNA-treated group of vehicle treated fibroblasts by two-way ANOVA.

### Effect of high-concentration trehalose on the expression of non-inflammatory SASP factor genes

The SASP is closely associated with positive and negative outcomes depending on cell types and contexts. Senescent cells also generate proinflammatory molecules and matrix metalloproteinases. Next, we investigated whether trehalose-induced cellular senescence in monolayer and organotypic cultures of human fibroblasts would lead to similarly dramatic changes in senescence factors induced by other stressors. Surprisingly, analysis of RNA-seq data revealed that SASP factor genes related to inflammation, such as *IL-6*, *IL-8*, and *IL-1B*, were not elevated, whereas several genes associated with senescence, such as *GDF15*, *MMP3*, and *TNFRSF10C*, were upregulated (*Table 1*). Overall, our data support the hypothesis that trehalose elicits the non-inflammatory SASP.

**Table 1.** Expression of SASP factor genes by highly concentrated trehalose. Analysis of RNA-seq data revealed that several genes associated with trehalose-induced premature senescence were upregulated.

| 2D Fibroblast |  |  |  | 3D Fibroblast |  |  |
| --- | --- | --- | --- | --- | --- | --- |
| Gene Symbol | Log<sub>2</sub> Fold Change | P value | q value | Log<sub>2</sub> Fold Change | P value | q value |
| CCL26 | 15.6887127 | 2.0027E-05 | 0.00044792 | 1.705299024 | 0.000739 | 0.008949 |
| CCL5 | 1.068414138 | 0.45589618 | 0.45288915 | -0.173989996 | 0.327876 | 0.492259 |
| CTSB | 0.971606011 | 0.06904278 | 0.1363981 | 0.335018077 | 0.000016 | 0.001276 |
| CXCL1 | 1.068140074 | 0.73874177 | 0.65627578 | -0.874653559 | 0.002441 | 0.019497 |
| CXCL5 | 0.94567579 | 0.75128719 | 0.66503826 | -0.777055659 | 0.001055 | 0.011256 |
| IL1B | 1.038811561 | 0.37390097 | 0.38579554 | -0.000435529 | 0.998081 | 0.929793 |
| IL8 | 0.749017012 | 0.40241364 | 0.4091766 | -1.863078226 | 0.000022 | 0.001431 |
| IL6 | 1.83106492 | 0.03268861 | 0.07368166 | 0.560710784 | 0.01188 | 0.058068 |
| IL6ST | 0.768246043 | 0.00028053 | 0.00211584 | -0.153220532 | 0.008595 | 0.046269 |
| IL6R | 0.961693449 | 0.60541119 | 0.56747153 | 0.340425901 | 0.022357 | 0.091428 |
| IL11 | 1.38408421 | 0.00127207 | 0.00602665 | -0.641457641 | 0.000063 | 0.002266 |
| GDF15 | 5.504128467 | 5.3168E-06 | 0.00021386 | 2.204582142 | 0.000023 | 0.001467 |
| ICAM3 | 1.10713058 | 0.17158772 | 0.26835683 | -0.04654121 | 0.784125 | 0.816121 |
| IGFBP2 | 0.816306772 | 0.21727315 | 0.32304459 | 0.153371435 | 0.499837 | 0.607437 |
| IGFBP4 | 0.676750709 | 1.1438E-05 | 0.0003231 | -0.096631441 | 0.019633 | 0.083265 |
| IGFBP6 | 0.599951743 | 2.1271E-05 | 0.0004627 | -0.440933601 | 0.000426 | 0.006395 |
| INHBA | 0.919449668 | 0.27440813 | 0.38579554 | 0.282262443 | 0.002374 | 0.019129 |
| KITLG | 1.193926054 | 0.00022606 | 0.00185082 | 0.153844064 | 0.005932 | 0.035706 |
| LIF | 0.733713608 | 0.00408197 | 0.01444751 | 0.198355932 | 0.053675 | 0.173018 |
| MMP1 | 1.107060554 | 0.00487661 | 0.01651 | 0.917704713 | 0.000062 | 0.00226 |
| MMP3 | 1.286485945 | 3.9542E-05 | 0.00063832 | 2.32521748 | <0.000001 | 0.000541 |
| MMP10 | 1.466808305 | 0.00679295 | 0.02126738 | 1.567801079 | 0.000048 | 0.002069 |
| MMP12 | 1.614463727 | 0.01064761 | 0.03018434 | 0.47100454 | 0.058762 | 0.185098 |
| MMP14 | 0.555567434 | 7.3144E-06 | 0.00025431 | 0.129684845 | 0.003357 | 0.024065 |
| PLAT | 2.985791621 | 9.1255E-06 | 0.00028893 | -0.25587261 | 0.025534 | 0.100313 |
| PLAU | 0.699589106 | 2.2605E-05 | 0.0004765 | -0.085737884 | 0.136164 | 0.323803 |
| PLAUR | 0.804912306 | 0.00707554 | 0.02193321 | -0.448530887 | 0.000271 | 0.004945 |
| SERPINB2 | 0.935304804 | 0.55097058 | 0.52692515 | -0.108924632 | 0.364082 | 0.492259 |
| TIMP1 | 0.899312334 | 0.00046492 | 0.00295382 | -0.13252159 | 0.01695 | 0.07489 |
| TNFRSF10C | 4.022071381 | 6.6203E-05 | 0.00085594 | 2.191704997 | 0.000001 | 0.000567 |
| STC1 | 0.872968807 | 0.08588855 | 0.16310923 | 0.268688045 | 0.008082 | 0.044319 |
| HMGB1 | 0.749005876 | 3.46E-05 | 0.00059345 | -0.672302379 | 0.000018 | 0.001331 |
| CALR | 0.844018275 | 3.6457E-06 | 0.00017579 | -0.358529449 | 0.000113 | 0.003023 |
| CD44 | 0.506002712 | 1.0211E-07 | 5.1976E-05 | -0.126432638 | 0.020103 | 0.084665 |
| S100A11 | 1.039862234 | 0.11871843 | 0.20483225 | 0.06647586 | 0.10953 | 0.293671 |
| LGALS3BP | 0.807693816 | 0.00822613 | 0.02470954 | -0.307984476 | 0.016317 | 0.07301 |
| VCAN | 0.720397578 | 0.00094683 | 0.00486125 | -0.47473911 | 0.000133 | 0.003325 |
| TNC | 3.665322378 | 1.0098E-07 | 5.1976E-05 | -0.100713946 | 0.023494 | 0.094632 |
| HSPA5 | 0.749125475 | 7.657E-06 | 0.00025961 | -0.618822695 | 0.000002 | 0.000603 |
| HSP90AB1 | 0.748200289 | 2.7946E-06 | 0.00015423 | -0.10722718 | 0.012148 | 0.059095 |
| HSPA8 | 1.358284902 | 1.1828E-05 | 0.00032903 | 0.014429089 | 0.628248 | 0.707638 |
| HSPA1A | 2.360615298 | 1.9545E-06 | 0.00012892 | -0.659152755 | 0.007447 | 0.041969 |
| HSP90AA1 | 1.105287269 | 0.00070951 | 0.0039587 | -0.063762606 | 0.018879 | 0.081054 |
| HSP90B1 | 0.645522626 | 7.4136E-07 | 0.00010197 | -0.874711901 | 0.000005 | 0.000797 |

**Table 2.** TaqMan primers and probe assay ID (Applied Biosystems) of the genes used in real-time PCR.

| Gene symbol | Assay ID | Gene Name |
| --- | --- | --- |
| EREG | Hs00914313_m1 | Epiregulin |
| AREG | Hs00950669_m1 | Amphiregulin |
| GAPDH | Hs02758991_g1 | Glyceraldehyde-3-Phosphate Dehydrogenase |
| ARG2 | Hs00982833_m1 | Arginase 2 |
| PGF | Hs00182176_m1 | Placental Growth Factor |
| VEGFA | Hs00900055_m1 | Vascular Endothelial Growth Factor A |
| DPT | Hs00355056_m1 | Dermapontin |
| CCL2 | Hs00234140_m1 | C-C Motif Chemokine Ligand 2 |
| SPP1 | Hs00959010_m1 | Secreted Phosphoprotein 1 |
| IL1RN | Hs00893626_m1 | Interleukin 1 Receptor Antagonist |
| ANGPT2 | Hs00169867_m1 | Angiopoietin 2 |
| AURKA | Hs00269212_m1 | Aurora Kinase A |
| AURKB | Hs00177782_m1 | Aurora Kinase B |
| AURKC | Hs00152930_m1 | Aurora Kinase C |
| CDKN1A | Hs00355782_m1 | Cyclin Dependent Kinase Inhibitor 1A |
| PLK1 | Hs00983227_m1 | Polo Like Kinase 1 |
| LMNB1 | Hs01059210_m1 | Lamin B1 |
| FGF2 | Hs00960934_m1 | Fibroblast Growth Factor 2 |
| MYBL2 | Hs00942540_m1 | MYB Proto-Oncogene Like 2 |
| UBE2C | Hs00153153_m1 | Ubiquitin Conjugating Enzyme E2 C |
| HIF1A | Hs00152153_m1 | Hypoxia Inducible Factor 1 Subunit Alpha |

### Dermal substitute with high-concentration trehalose-treated fibroblasts enhances wound closure and capillary formation

Since trehalose treatment upregulated wound healing-related genes in the 3D culture system, 6 mm × 6 mm full skin thickness excisional wounds were made on the dorsum of nude mice (BALB/cAJcl-nu). The dermal substitutes composed of collagen and fibroblasts with or without trehalose (100 mg/ml) were further transplanted to test whether trehalose-treated fibroblasts in the dermal substitute accelerate wound closure *in vivo*. In the macromorphological analysis, we detected a significant tendency toward promoted healing in the high-concentration trehalose-treated group compared with the control group (*Figure 8, A and B*). Induction of neoangiogenesis was observed 7 days after transplantation of the dermal substitute with high-concentration trehalose (*Figure 8C*). Furthermore, statistical analysis demonstrated a significant difference in the narrower wound opening of the trehalose-treated dermal substitute group as detected histologically by hematoxylin and eosin staining 7 days after transplantation (*Figure 8, D and E*). We also stained the tissue with antibodies against CD31. Compared with that in the vehicle-treated group, a significantly greater number of vessels in the wounds transplanted with the trehalose-treated dermal substitute stained positive for CD31 (*Figure 8, F and G*). Therefore, the dermal substitute with trehalose-treated fibroblasts accelerated wound healing by promoting capillary formation *in vivo*.

**Figure 8.**
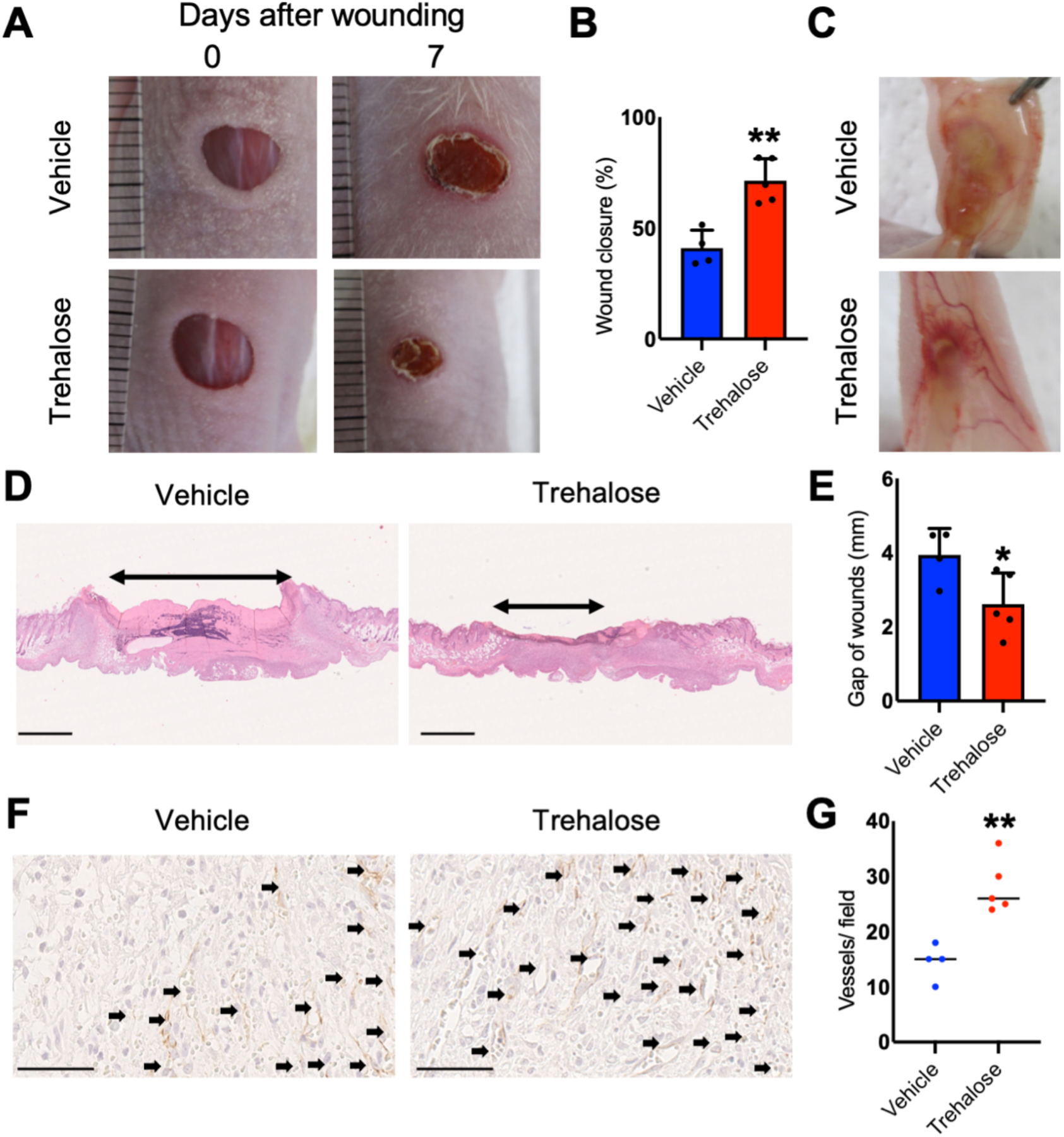
Graft of dermal substitute with highly concentrated trehalose-treated fibroblasts on nude mice accelerates murine wound closure and angiogenesis. (**A**) Representative photographs of the wound area on the recipient nude mice as indicated at 7 days after transplantation. (**B**) Wound closure was quantified and presented as % wound closure; % = the percentage of the initial wound area size at day 7 when comparing the trehalose-LSE group (red band) to the control group (blue band). (**C**) Representative photographs of angiogenesis induced by the grafts. (**D**) Images of the H&E-stained tissue sections from the wound sites at day 7. Bars = 1 mm. (**E**) Quantitative analysis of gap of the wounds in comparison between the trehalose-LSE group (red band) and control group (blue band). (**F**) Images of CD31 immunostaining on the day 7 wounds. Bars = 50 μm (**G**) Quantitative analysis of CD31 positive vessels per field. Data are expressed as means ± SD. A subsequent statistical analysis was performed with Student *t*-test. *: *P* < 0.05, **: *P* < 0.01, with four mice in the control group, five in the trehalose-LSE group, and one tissue section from each mouse. Data are representative of three independent experiments.

## DISCUSSION

The challenge to engineer cultured epidermal autografts for the life-saving treatment of patients with extensive, full-thickness burns was accomplished using the method of keratinocyte-cultivation described by Rheinwald and Green (O’Connor et al., 1981, Rheinwald and Green, 1975). However, their method requires the use of a feeder layer of lethally irradiated mouse 3T3 cells and serum. Therefore, the regulatory issues have necessitated the use of xenotransplantation and development of cultivation technology. Moreover, epidermal grafts without the dermis are less resistant to trauma and more prone to post-transplantation contracture, leading to the poor functional and cosmetic outcomes. An autograft full-thickness LSE can not be used to treat patients with burns due to the time required for preparation, despite the advances in the methods for rapid *ex vivo* expansion. In this report, we demonstrated a breakthrough for the new techniques for the rapid development of LSE using the effect of highly concentrated trehalose added to collagen gel, even as a pretreatment, which induces the transient beneficial SASP in human fibroblasts via CDKN1A/p21 and modulates the capacity to accelerate the proliferation of keratinocytes in the epidermal layer of LSE. Furthermore, the trehalose-treated skin equivalents promoted wound repair with an angiogenic effect *in vivo* (*Figure 9*).

**Figure 9.**
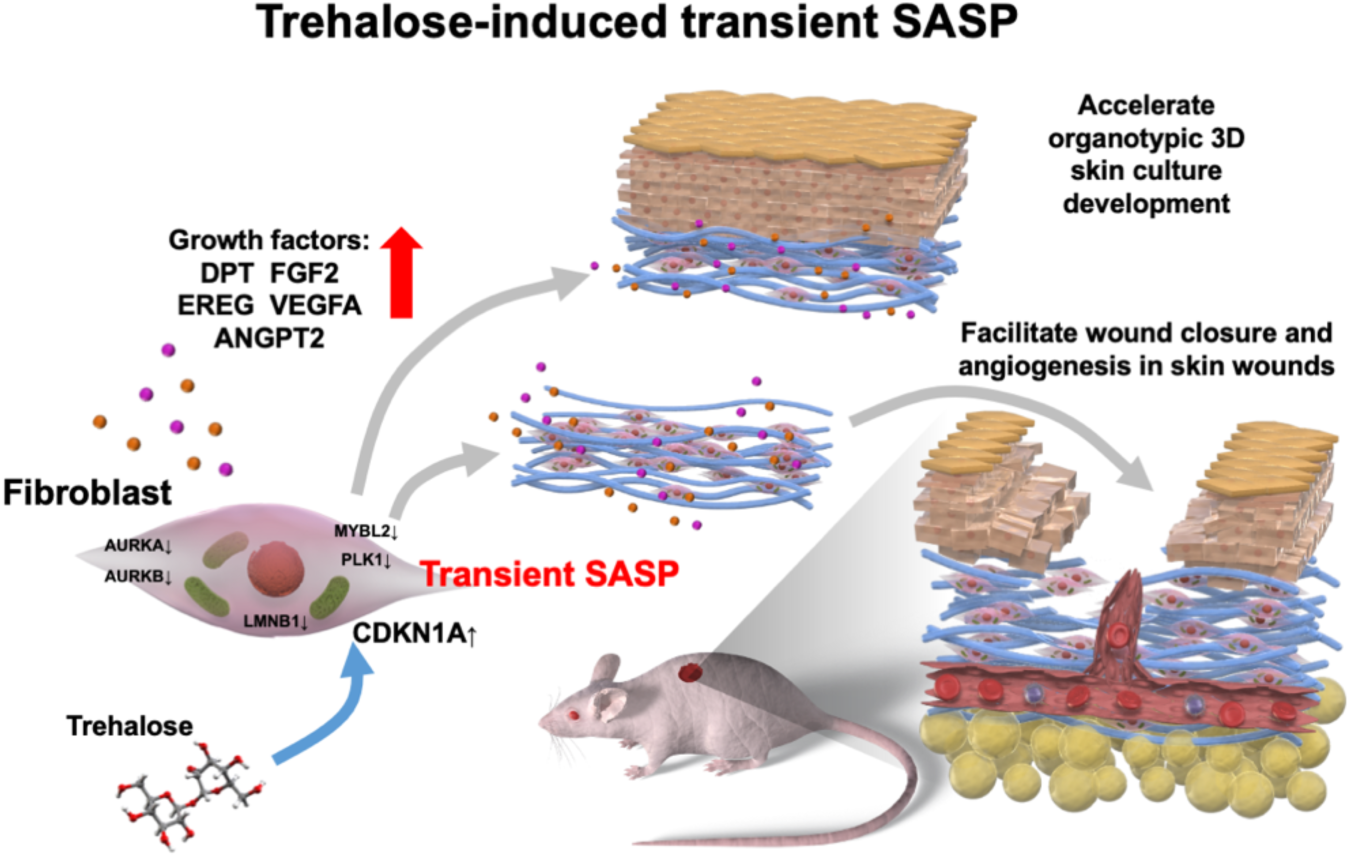
Graphical abstract: Novel trehalose-induced transient SASP accelerates organotypic skin culture generation and facilitates cutaneous wound closure.

These observations elucidate important physiologic roles for fibroblasts in the construction of LSE *in vitro* and in wound repair *in vivo*. Our 3D culture system confirmed the role of trehalose-treated fibroblasts in keratinocyte proliferation. Somewhat surprisingly, we observed the upregulation of growth factors such as DPT, FGF2, EREG, VEGF, and ANGPT2 as growth factors in the highly concentrated trehalose-induced SASP. Importantly, we discovered that this effect of trehalose was transient and occurred only at an early time point, because RNA-seq analysis revealed similar gene expression profiles for trehalose-treated and the vehicle-treated keratinocytes and fibroblasts in the final LSE preparations. Furthermore, we found that trehalose-induced SASP was relatively non-inflammatory compared with other stress-induced SASPs. Therefore, trehalose-induced transient non-inflammatory SASP of human fibroblasts is a novel approach for accelerating keratinocyte proliferation in LSE and wound healing *in vivo*. We propose this trehalose-induced SASP to be named “trehalose-induced senescence-associated secretory phenotype (TISASP).” These composite grafts associating autologous keratinocytes with fibroblasts may have a major impact on chronic wound therapy. We observed that the addition of sucrose of the same concentration (100 mg/ml) in the medium induced cell death in human dermal fibroblasts, suggesting that TISASP is not due to the stress of the disaccharide-induced osmotic pressure.

The use of RNA-seq technology enables an unbiased, sensitive method for investigating the transcriptome of LSEs, 3D fibroblasts, and the 2D monolayer under trehalose treatment. We identified significantly and differentially expressed genes overlapping between the two data sets for 2D and 3D fibroblasts. RNA-seq analysis data and the IPA of the differentially expressed genes revealed a potential key role for the CDKN1A pathway for transient G2 arrest based on the discovery of the upregulation of other growth factors including DPT. The *CDKN1A* gene represents a major target of the p53 activity, and its product, p21, is the major regulator of the cellular stress response (Warfel and El-Deiry, 2013). The capacity of p21 for cell cycle arrest depends upon its nuclear localization (Wu et al., 2011). Our immunocytochemistry results revealed that p21 levels increased mainly in the nucleus after trehalose treatment. Additionally, the RNA-seq data revealed inhibition of AURKA and AURKB in the fibroblasts treated with trehalose. The Aurora kinases, including AURKA and AURKB are highly conserved serine/threonine kinases essential for the control of mitosis. AURKB indirectly represses the expression of CDKN1A at the transcriptional level (Trakala et al., 2013). Thus, downregulation of AURKB leads to the induction of p21. Treatment of cultured multiple myeloma cells with MLN8237, which is a small-molecule and Aurora-A kinase inhibitor that inhibits Aurora-A gene expression by siRNA, results in G2/M arrest and senescence *in vitro* (Dutta-Simmons et al., 2009). P21 inhibits AURKA by regulating E2F3 (Wu et al., 2012). Hence, under the stress of high-concentration trehalose, induced transcription of *CDKN1A* inhibited AURKA, which led to the G2/M blockade (CDK1-induced mitotic entry). Furthermore, upstream analysis of RNA-seq data revealed the involvement of ZBTB17, which binds to the *AURKA* promoter and is assumed to be associated with transcriptional factors that induce *AURKA* downregulation following topoisomerase I inhibition (Courapied et al., 2010).

DeBosch *et al*. reported that activity of glucose transporters at the plasma membrane (SLC2A1: GLUT1, SLC2A2: GLUT2, SLC2A3: GLUT3, SLC2A4: GLUT4, and SLC2A8: GLUT8), which are expressed in fibroblasts (Longo et al., 1990), is inhibited by 100 mM trehalose (DeBosch et al., 2016). We treated the fibroblasts with 29.2 (10 mg/ml) to 292 mM (100 mg/ml) trehalose and observed a dose-dependent effect in our system. Upon glucose restriction in the medium (4.4 mM), yeast cells underwent transient cell cycle arrest at the G2 phase, which is dependent on the Wee1 tyrosine kinase (Masuda et al., 2016). We identified a significant decrease of *WEE1* mRNA levels in trehalose-treated 2D and 3D fibroblasts compared with that of the control in our RNA-seq data. Therefore, trehalose-mediated inhibition of glucose absorption in human dermal fibroblasts might lead to transient Wee1-independent G2 cell cycle arrest in our system. We also identified increased mRNA levels of *HIF1A* after trehalose treatment in the cultured monolayer and 3D fibroblasts. Interestingly, the levels of reactive oxygen species (ROS) in both the mitochondria and the cytosol increased upon glucose withdrawal, although reduced to the background level upon glucose re-feeding in human newborn foreskin fibroblasts (Song and Hwang, 2018). There was significantly higher superoxide radical generation in those fibroblasts treated with trehalose (*Figure 5-figure supplement 2*). The growing body of evidence suggests that ROS induces hypoxia-inducible factor 1-α (HIF1α) via MAPK, ERK, and PI3K/AKT pathways (Movafagh et al., 2015). HIF1α negatively regulates AURKA in breast cancer cell lines under hypoxic conditions (Fanale et al., 2013) and is involved in *CDKN1A* transcription in murine embryonic fibroblasts (Goda et al., 2003). Hence, elevated ROS levels after glucose transport inhibition in trehalose-treated fibroblasts induced HIF1α and could be involved in the subsequent transcription of *CDKN1A*, thus leading to a G2/M blockade in our system.

The role of CDKN1A is likely confined to the induction of senescence and cells can resume cycling upon resolution of stress (Childs et al., 2015). In embryonic development, CDKN1A induction leads to SASP factor expressions, like FGFs, which stimulate cell proliferation and tissue formation (Da Silva-Alvarez et al., 2019). Importantly, we found highly concentrated trehalose can markedly increase CDKN1A/ p21 expression, which is required for a striking upregulation of FGF2 and other growth factors. Demaria *et al*. analyzed p16^INK4a^/CDKN1A double knockout mice, and found that the wound healing of knockout mice was impaired compared with their wild-type controls, thus indicating that the presence of senescent cells facilitates skin wound healing, and their absence significantly suppresses wound closure (Demaria et al., 2014). Thus, transient senescence is critical for effective cutaneous wound healing. We hypothesize that trehalose treatment can induce a transient and beneficial senescent phenotype of the human fibroblasts via CDKN1A/p21 for optimal wound healing.

Previous studies demonstrated that epidermal growth factor (EGF) family members—transforming growth factor (TGF)-α, heparin-binding (HB)-EGF, and EREG—act as autocrine growth factors for normal human keratinocytes (Shirakata, 2010). We also observed dramatic upregulation of *EREG* after trehalose treatment. EREG is upregulated in the psoriatic epidermis and was initially purified from the mouse NIH-3T3 (Shirakata, 2010). We also demonstrated that trehalose treatment induced a marked and significant increase of mRNA and protein levels of DPT, which is a 22-kDa matrix protein for the interaction with TGFβ1, fibronectin, and decorin. DPT dose-dependently promotes keratinocyte migration in wound repair (Krishnaswamy and Korrapati, 2014). Furthermore, trehalose attenuates protein aggregation and maintains polypeptide chains in a partially folded state for the refolding by cellular chaperones (Singer and Lindquist, 1998). Therefore, we speculate that trehalose plays a role in inducing the secretion of growth factors from fibroblasts and facilitates the functions of these overexpressed growth factors as chaperones for keratinocyte proliferation on collagen gels.

Although CDKN1A is a vital senescence marker, it is induced during transient cell cycle arrest, thus must be used in combination with other markers (Herranz and Gil, 2018). A recently emerging candidate marker that seems to play a role in attenuating senescence is MYBL2, which is a transcription factor of the MYB family (Musa et al., 2017). The p53–p21 pathway suppresses MYBL2 expression as a stress response (Fischer et al., 2016). Our RNA-seq results demonstrated that downregulation of *MYBL2* and target genes transactivated by MYBL2, such as *AURKA*, *CCNA2*, *CCNB1*, *CDK1*, *PLK1*, and *TOP2A*, are present in human fibroblasts following trehalose treatment, which indicates that MYBL2 may participate in the senescence phenotype of human fibroblasts after trehalose treatment. Furthermore, reduced *LMNB1* mRNA expression strongly predicts the senescence phenotype. LMNB1 expression decreased in human fibroblasts after trehalose treatment. Thus, LMNB1 can be a marker for trehalose-induced senescence.

Pertinent roles for transient senescence in tissue injury have been identified during wound repair: PDGFA-enriched SASP (Demaria et al., 2014). SASP factors include CCL2, which is a chemokine required for the chemotaxis of macrophages and monocytes during angiogenesis in wound repair. Recently, Whelan *et al*. demonstrated a novel role of mesenchymal stromal cell-derived CCL2 in accelerated wound closure (Whelan et al., 2020). In addition, the absence of the IL-1 receptor antagonist (IL1RN) impaired wound healing along with aberrant NF-κB activation and reciprocal suppression of the TGF-β pathway (Ishida et al., 2006). Placental growth factor (PGF) encodes a growth factor that is homologous to the vascular endothelial growth factor and is a potent angiogenic/permeability factor during wound repair (Failla et al., 2000). Osteopontin (OPN) is a glycoprotein that is encoded by secreted phosphoprotein 1 gene (*SPP1*), and analysis of *OPN* null mutant mice indicated the *in vivo* role of OPN in structural remodeling and resolution of dermal wounds (Liaw et al., 1998). Angiopoietin 2 encoded by the *ANGPT2* gene acts as a Tie-2 antagonist (Maisonpierre et al., 1997), thus increasing sensitivity to other proangiogenic factors such as VEGF (Holash et al., 1999). We demonstrated significant upregulation of these wound healing-related genes, such as *EREG*, *CCL2*, *IL1RN*, *PGF*, *SPP1*, *VEGFA*, *DPT*, *FGF2*, *ARG2*, and *ANGPT2*, using RNA-seq and qPCR mRNA expression analysis in trehalose-treated 2D and 3D fibroblasts. Furthermore, we found that transplantation of a dermal substitute prepared with collagen gel and fibroblasts treated with trehalose promoted wound healing and capillary formation *in vivo* compared with wound recovery of the vehicle-treated control. Considered together, trehalose-induced angiopoietin 2 may act in concert with these SASP factors, such as VEGFA, to stimulate angiogenesis in the wounds. Although the results are limited to the murine model, we provide evidence that induction of the transient non-inflammatory senescence-like phenotype by trehalose in fibroblasts is beneficial for healing skin wounds. In a p53-induced senescence model, cooperation between p21 and Akt was required for inducing the cellular senescence phenotype and cell cycle arrest (Kim et al., 2017). Interestingly, we here show that high-concentration trehalose activates ERK1/2 and AKT in human fibroblasts. By contrast, Wu *et al*. reported intraperitoneally-injected trehalose promotes the survival of rat skin flaps and angiogenesis by autophagy enhancement due to inhibition of Akt (Wu et al., 2019).Thus, high-concentration trehalose’s mechanism of action is different from that previously known. Moreover, PGF and VEGFA accelerate diabetic wound healing, so these gene transfers to diabetic wounds have received increasing attention (Cianfarani et al., 2006, Sun et al., 2018). Topical application of CCL2 can accelerate cutaneous wound healing in mice with diabetes by promoting neovascularization (Ishida et al., 2019). Therefore, future senescence-targeted therapies with trehalose should be reserved for the treatment of chronic wounds of human diabetic patients.

In conclusion, this study demonstrated that highly concentrated trehalose induces transient SASP in fibroblasts, and revealed that trehalose-induced cell cycle arrest and growth factor secretion via CDKN1A/p21 are beneficial for keratinocyte proliferation in LSE construction *in vitro* and capillary formation and wound closure in the repair process *in vivo*. These data suggest a new therapeutic approach for altering wound responses by applying trehalose-treated fibroblasts for accelerating wound repair. This could provide the foundation of a new therapy to treat not only genodermatoses such as epidermolysis bullosa but also chronic diabetic and venous ulcers. Therefore, we believe these findings should promote future studies on the effect of trehalose in modulating fibroblast functions for LSE construction and subsequent wound therapy.

## MATERIALS AND METHODS

### Chemicals and reagents

Trehalose (containing >98.0% trehalose dihydrate, Hayashibara, Okayama, Japan), oligo-hyaluronan (hyaluronan oligosaccharide 4mer, CSR-11006, Cosmo Bio, Tokyo, Japan), and biotin-labeled hyaluronic acid-binding protein (HKD, Sapporo, Japan) were purchased.

### Cell culture

Normal human epidermal keratinocytes were isolated from normal human skin and cultured under serum-free conditions, following a previously described method (Shirakata et al., 2003, Shirakata et al., 2004). The cells were used for LSE cultures in their fourth passage. Fibroblasts were isolated from normal human skin and cultured in Dulbecco’s modified Eagle medium (DMEM) (Thermo Fisher Scientific) supplemented with 10% fetal calf serum, and fifth-passage cells were used to construct the LSEs. All procedures that involved human subjects received prior approval from the Ethics Committee of Ehime University School of Medicine, Toon, Ehime, Japan, and all subjects provided written informed consent.

### Preparation of cultured skin equivalents with or without trehalose

The method used for LSE preparation was described previously (Yang et al., 2005). Briefly, a collagen gel was prepared by mixing six volumes of ice-cold porcine collagen type I solution (Nitta Gelatin, Osaka, Japan) with one volume of 8 × DMEM, 10 volumes of 1 × DMEM supplemented with 20% FBS, and one volume of 0.1 N NaOH, of which 1 ml was added to each culture insert (Transwell, 3-µm membrane pore, Corning, Corning, NY) in a six-well culture plate (Corning). Following polymerization of the gel in the inserts at 37°C, two volumes of fibroblast suspension solution 5 × 10^5^ cells/ml in 1 × DMEM supplemented with 10% FBS were added to eight volumes of the collagen solution (thus, the final collagen concentration was 0.8 mg/ml). Then, 3.5 ml of the fibroblast-containing collagen solution was applied to each insert. When the fibroblast-containing gel polymerized, DMEM supplemented with 10% FBS and ascorbic acid (final concentration 50 ng/ml) was added with or without trehalose (in three concentrations: 10, 30, and 100 mg/ml). The gel was submerged in culture for 5 days until the fibroblasts contracted the gel.

A larger LSE was constructed following the same method as previously described except using a larger culture insert (Transwell, 75-mm diameter, 3-µm membrane pore, Corning), thus utilizing proportionally more fibroblasts. A rubber ring (8-mm interior diameter) was covered over the fibroblast-containing gel to stabilize it within the large-scale LSE. In the hole of the ring, 6 × 10^5^ keratinocytes in 30 µl MCDB 153 type II were seeded. The keratinocytes were submerged in culture for 2 days. When the keratinocytes reached confluence, the LSE was lifted to the air–liquid interface and a cornification medium (a 1:1 mixture of Ham’s F-12 and DMEM supplemented with 2% FBS and other supplements, as described previously (Yang et al., 2005)) was added. The medium was changed every other day.

To construct the conventional LSE, keratinocytes were seeded onto the contracted gel and then submerged and airlifted as described above, except without the ring. The seeding cell density was adjusted using rubber rings. Both LSE types were harvested 7 or 14 days after airlifting. For hematoxylin and eosin staining, the LSE was fixed in 10% formalin and embedded in paraffin. For immunohistochemical staining, the LSE was snap frozen in an OCT compound. We performed more than five experiments and obtained similar results. A representative experiment depicted in the Figure 1. In comparative studies, keratinocytes and fibroblasts from the same donor were used.

### Transplanting cultured 3D dermal sheets

The animal grafting protocol was approved by the Ethics Committee of Ehime University School of Medicine. Ten-week-old female BALB/cAJc1-nu nude mice (CLEA Japan, Tokyo, Japan) were anesthetized by isoflurane inhalation. Full-thickness wounds were created on the skin of the backs of each mouse using a 6-mm skin biopsy punch. The fibroblast-containing collagen gels were prepared with vehicle or trehalose (100 mg/ml) and submerged in culture for 5 days, and the dermal substitutes (1 day after airlift) were grafted onto the wounds, which were covered with transparent films. Seven days after transplantation, the grafts were harvested. One part of each graft was paraffin-embedded and sectioned at 6 µm, from which hematoxylin and eosin staining was prepared. Some sections were de-paraffinized and blocked for endogenous peroxidase activity, and then blood vessels were stained with rat antibody against CD31 (at 1:20 dilution, dianova GmbH, Hamburg, Germany), according to the manufacturer’s instructions for the ImmPRESS^TM^ reagent kit (Vector Laboratories, Burlingame, CA). The sections were counterstained with hematoxylin for cell nuclei. We performed at least three independent studies and confirmed similar results. A representative experiment is shown in the figures.

### Whole transcriptome analysis with RNA-seq

Total RNA was extracted from the fibroblasts or LSE using the RNeasy Mini Kit (Qiagen, Hilden, Germany), and mRNA was purified with oligo dT beads (NEBNext Poly (A) mRNA magnet Isolation Module, New England Biolabs, NEB, Ipswich, MA). The procedure of the complementary DNA (cDNA) libraries was carried out with NEBNext Ultra II RNA library Prep kit (NEB) and NEBNextplex Oligos for Illumina following a previously described method (Kohno et al., 2020). Briefly, mRNA was incubated in NEBNext First Strand Synthesis Reaction Buffer at 94°C for 15 min in the presence of NEBNext Random Primers, and reverse transcription was carried out with NEBNext Strand Synthesis Enzyme Mix. The index sequences were inserted to the fragments with PCR amplification. The libraries were added in equal molecular amounts and were sequenced on an Illumina Next-seq DNA sequencer with a 75-bp pair-end cycle sequencing kit (Illumina, San Diego, CA). The detected reads were analyzed using CLC Genomics Workbench software (ver.8.01, Qiagen). The pathway for the detected genes was analyzed using IPA (Qiagen).

### Histology and immunohistochemical staining

Paraffin-embedded LSE samples were sectioned at 6 µm and stained with hematoxylin and eosin (H&E) or alcian blue (pH 2.5). For immunohistochemical staining, ImmPRESS^TM^ reagent kit (Vector Laboratories) was used according to the manufacturer’s instructions. Frozen sections (7 µm) were first incubated with 0.3% hydrogen peroxide for 30 min to remove endogenous peroxidase activity and then incubated with primary antibodies at appropriate dilutions overnight at 4°C. The antibodies used in this study were n1584 for α-SMA (Agilent technologies, Santa Clara, CA) and NCL-Ki67-MM1 for Ki67 (LeicaBiosystems, Buffalo Grove, IL). The sections were incubated with enzyme-conjugated secondary antibodies for 30 min at room temperature and then with the staining substrate. To determine if hyaluronan accumulates in LSEs, staining was carried out using biotinylated-hyaluronic acid-binding protein (Cosmo Bio). To detect elastic fibers in the tissue, EVG staining was performed using standard histological dyes for EVG staining (Muto Pure Chemicals, Tokyo, Japan). Images were obtained using Nikon ECLIPSE E600 microscope coupled with Nikon DS-Ri1camera (Nikon, Tokyo, Japan).

### Evaluating the epidermal spreading potential of trehalose-treated LSEs

Rubber rings with an inner diameter of 8 mm were put on gels with or without trehalose in the gel (10, 30, and 100 mg/ml) and 6 × 10^5^ keratinocytes in 30 µl MCDB 153 II medium were seeded into each ring hole. When the keratinocytes reached confluence, the LSEs were lifted to air–liquid surface, and the rubber rings were removed. At 14 days after airlifting, the epidermal size was measured using computer-assisted morphometric analysis. The epidermal sizes of the conventional LSEs and trehalose-treated LSEs were compared statistically using Student’s *t*-test.

### SA-βgal assay, p21 and dihydroethidium (DHE) immunocytochemistry

SA-βAgal assay was performed by seeding fibroblasts onto eight-well chamber slides and treating with trehalose or vehicle. Cells were also treated with adenovirus vectors that encode lacZ (Ax LacZ), following a previously described method (Tokumaru et al., 2005). Cells were fixed and stained with the Senescence Detection Kit (Cell Signaling Technology, Danvers, MA), following the manufacturer’s instructions. The results were observed under a microscope. For p21 immunocytochemistry, the treated fibroblasts were fixed with 4% paraformaldehyde/phosphate-buffered saline (PBS) for 30 min at room temperature, permeabilized with 0.5% Triton X-100/PBS for 15 min, incubated with antibodies raised against p21 #29475 (Cell Signaling Technology) overnight at 4°C after blocking with blocking solution (S3022, Dako) for 30 min, and then incubated with an Alexa Fluor 488-conjugated secondary antibody and DAPI (Thermo Fisher Scientific). The cells were mounted with VECTASHIELD (Vector Laboratories). Fluorescence was observed with a fluorescence microscope (Nikon) and analyzed by ImageJ (National Institutes of Health). For evaluating the superoxide production, the staining was done with DHE (ab145360, Abcam, Cambridge, UK) in dark and fluorescence images were taken using fluorescence microscope BZ9000 (Keyence, Osaka, Japan) with fluorescent filter OP-66838 (excitation 560/30 nm and emission 630/60 nm). Fluorescent signals were quantified using ImageJ (National Institutes of Health).

### Cell death assays

Cell viability was measured using a Cell Counting Kit-8 assay (Enzo life sciences, Farmingdale, NY) following the manufacturer’s instructions. Optical density was measured at 450 nm and was normalized to the corresponding stimulation control.

### Cell cycle analysis of monolayer fibroblasts and dermal cells in LSEs

Single-cell preparations from the monolayer fibroblasts or the dermal side of LSE were carried out. The dermis was removed from the epidermis of the LSE cells. The dermis was further digested with collagenase XI and hyaluronidase (both from Sigma-Aldrich) for 120 min followed by fluorescence-activated cell sorting (FACS) analysis with propidium iodide (BioLegend, San Diego, CA) according to the propidium iodide cell cycle staining protocol.

### RNA preparation and determination of mRNA expression by quantitative RT-PCR

Total RNA was isolated by using the RNeasy Mini Kit (Qiagen), and real-time PCR was used to determine the mRNA abundance, as described previously (Dai et al., 2011). TaqMan™ Gene Expression Assays (Thermo Fisher Scientific) were used to analyze the expressions (Table S2)*. GAPDH* mRNA was used as an internal control. Target gene mRNA expression was calculated relative to *GAPDH* mRNA, and all data are presented as normalized data compared to each control (mean of control cells or tissues).

### Small interfering RNA

Silencer validated siRNA CDKN1A (AM51331, Thermo Fisher Scientific) was used for silencing CDKN1A, and the Silencer negative control siRNA (AM4611, Thermo Fisher Scientific) was used as control. Fibroblasts were transfected with siRNA using Lipofectamine RNAiMAX Transfection Reagent (Thermo Fisher Scientific) according to the manufacturer’s instructions. The cells were allowed to stabilized for 24 hours before trehalose or vehicle treatment.

### Western blotting analysis

Following stimulation, total cell extracts were collected at the indicated times. To detect the protein levels, cell lysates were separated by SDS–polyacrylamide gel electrophoresis and transferred to polyvinylidene difluoride membranes. Analyses were performed using Amersham ECL Prime Western Blotting Detection Reagent (RPN2232) (GE Healthcare Life Sciences, Chicago, IL), and then the membranes were scanned using Image Quant LAS4010 (GE Healthcare Life Sciences). We obtained the primary antibodies for ERK (#9102), phospho-ERK (#4370), AKT (#9272), phospho-AKT (#9271), p21 waf/cipl (12p1) (#29475) (Cell Signaling Technology), LaminB (C-20) (catalog sc-6216) (Santa Cruz, Dallas, TX), DPT (AF4629) (R&D systems, Minneapolis, MN), and β-actin (#ab6276)(Abcam).

### DPT ELISA

DPT in the cell culture supernatants were measured. A Human DPT ELISA Kit (Abcam) was used to measure DPT in LSE, according to the manufacturer’s procedures.

### Statistics

Statistical analysis was performed using a two-tailed Student’s *t*-test, one- or two-way analysis of variance with Prism software (version 9; GraphPad Software, San Diego, CA). Results are expressed as the mean ± standard deviation (SD). A *P*-value of <0.05 was considered significant.

### Study approval

All animal procedures performed in this study were reviewed and approved by the Ehime University Institutional Animal Care and Use Committee. The experiments were conducted in accordance with the NIH guidelines for care and use of animals and the recommendations of International Association for the Study of Pain.

## Acknowledgments

This work was supported by TR SPRINT Stage-A Seeds (A209 and A054) from Japan Agency for Medical Research and Development (AMED). The authors would like to thank E. Tan and Dr. K. Kameda from Ehime University for performing the qPCR and flow cytometry experiments.

## Data and materials availability

All data are available in the paper or the supplementary materials. RNA sequence data are submitted to GEO under accession number GSE184892.

The following data sets were generated.

Jun Muto, Shinji Fukuda, Kenji Watanabe, Xiuju Dai, Teruko Tsuda, Takeshi Kiyoi, Hideki Mori, Ken Shiraishi, Masamoto Murakami, Shigeki Higashiyama, Yoichi Mizukami, Koji Sayama (2021) NCBI Gene Expression Omnibus ID GSE 184892.

Highly concentrated trehalose induces transient senescence-associated secretory phenotype in fibroblasts via CDKN1A/p21

https://www.ncbi.nlm.nih.gov/geo/query/acc.cgi?acc=GSE184892

## Definitions of abbreviations

ANGPT2: angiopoietin-2
α-SMA: α-smooth muscle actin
AURKA: Aurora kinase A
AURKB: Aurora kinase B
CDKN1A: cyclin-dependent kinase inhibitor 1A
DPT: dermapontin
EREG: epiregulin
FGF: fibroblast growth factor
GM-CSF: granulocyte macrophage colony-stimulating factor
HABP: hyaluronan-binding protein
HIF1α: hypoxia-inducible factor 1-α
HA: hyaluronan
IGF: insulin-like growth factor
IL1RN: IL-1 receptor antagonist
IPA: Ingenuity Pathway Analysis
LMNB1: Lamin B1
LSE: living skin equivalent
MYBL2: Myb proto-oncogene like 2
OPN: Osteopontin
PCA: Principal component analysis
PDGFA: platelet-derived growth factor-A
PGF: placental growth factor
PLK1: Polo-like kinase 1
qPCR: quantitative PCR
RNA-seq: RNA-sequencing
ROS: reactive oxygen species
SA β-gal: senescence-associated β-galactosidase
SASP: senescence-associated secretory phenotype
TGF: transforming growth factor
VEGF: vascular endothelial growth factor.

## Supplemental Information

**Figure 1-figure supplement 1:**
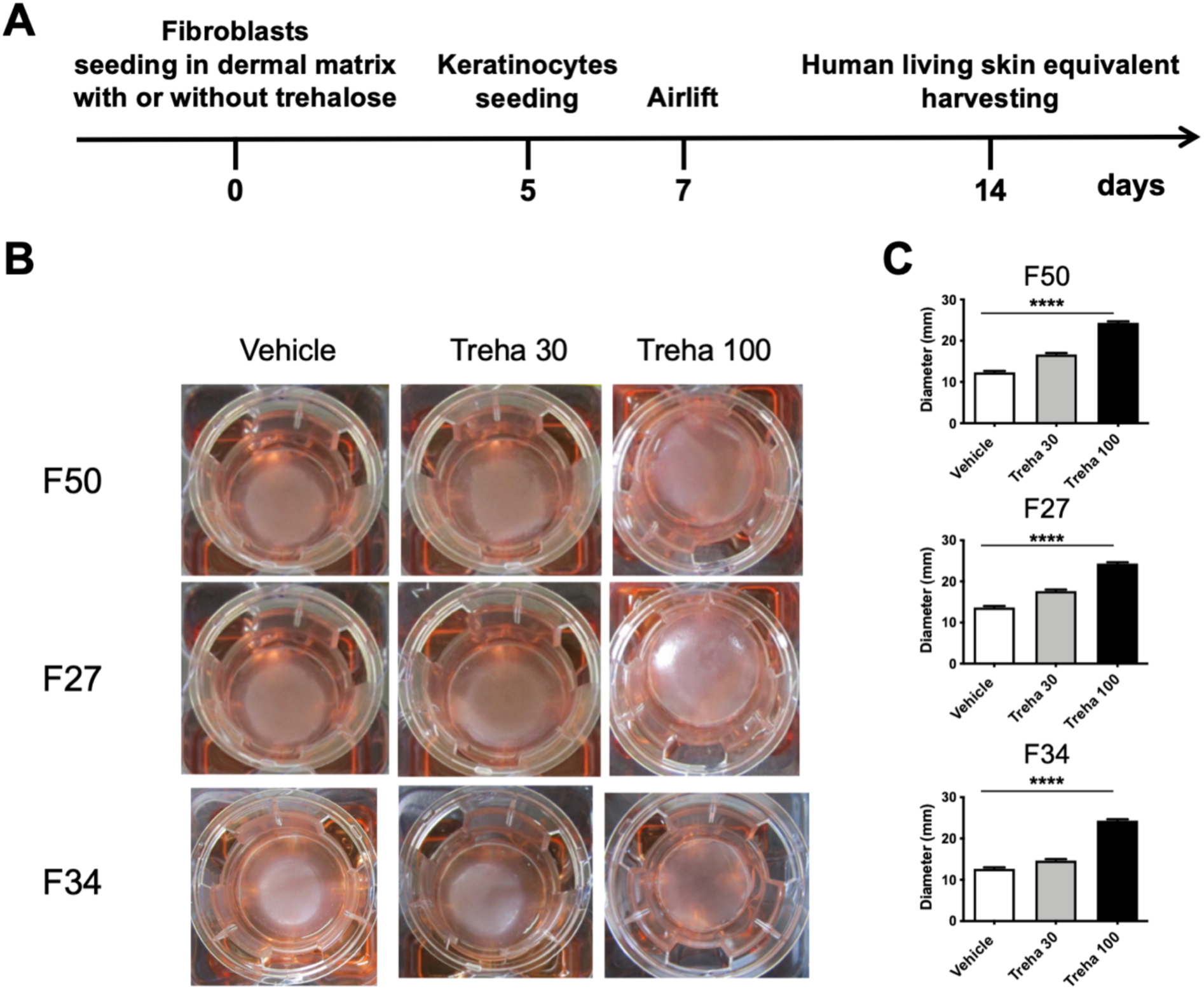
Novel effect of highly concentrated trehalose in the preparation of living skin equivalents. (**A**) A schematic for the preparation of cultured skin equivalents. (**B**) Macroscopic pictures of LSEs with and without trehalose (30 and 100 mg/ml) added in the collagen gel. The gel was prepared in the Transwell-COL with 24-mm insert in a six-well culture plate after 1-week of airlifting at 37°C. (**C**) Diameters of LSEs with and without trehalose added in the collagen gel after 1-week of airlifting at 37°C. Data are expressed as means ± SD for three LSEs. ****: *P* < 0.0001 versus vehicle control groups in Student *t*-test.

**Figure 1-figure supplement 2:**
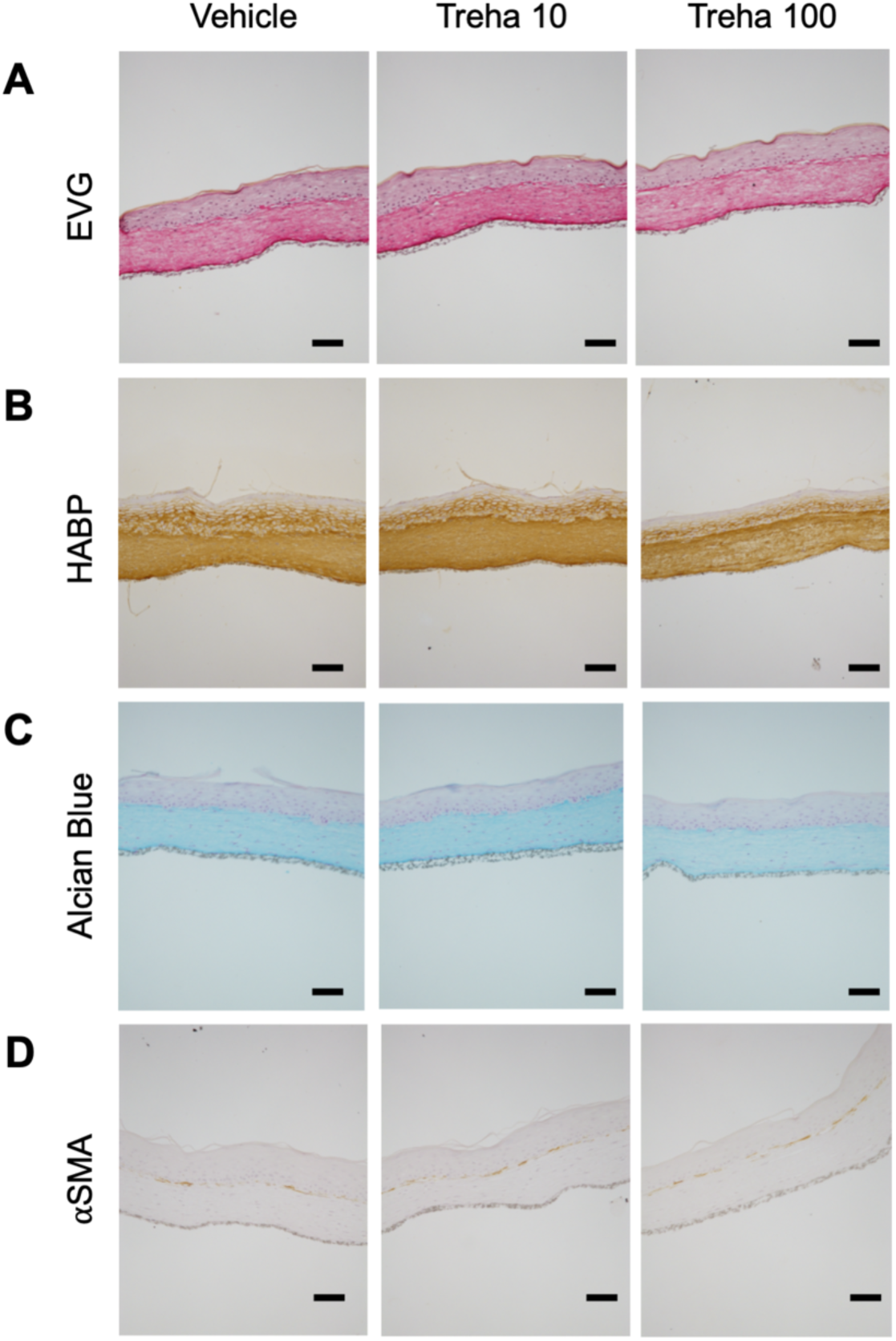
Immunostaining of living skin equivalents prepared with or without trehalose (10 and 100 mg/ml). Paraffin-embedded sections of LSEs were sectioned and subjected to immunohistochemistry for Elastica van Gieson staining (**A**), hyaluronan-binding protein (HABP) staining (**B**), Alcian blue staining, PH 2.5 (**C**), α-SMA staining (**D**). Scale bar = 50 μm.

**Figure 1-figure supplement 3:**
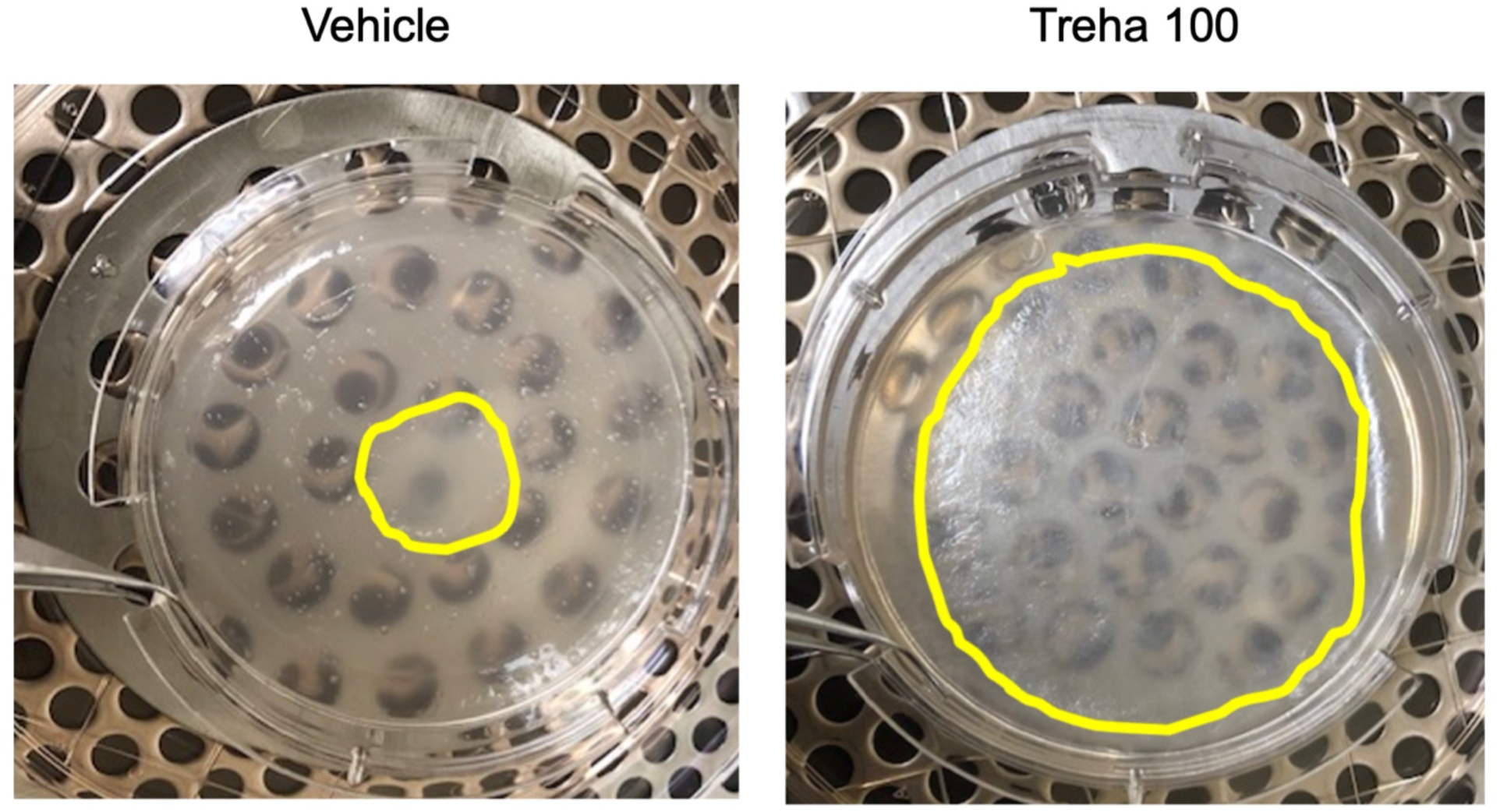
Novel effect of trehalose in the preparation of living skin equivalents. Representative picture of LSEs with or without trehalose (100 mg/ml) added in the collagen gel, which was prepared in a 100-mm dish after 2-weeks of airlifting at 37°C. Data are representative of two independent experiments.

**Figure 2-figure supplement 1:**
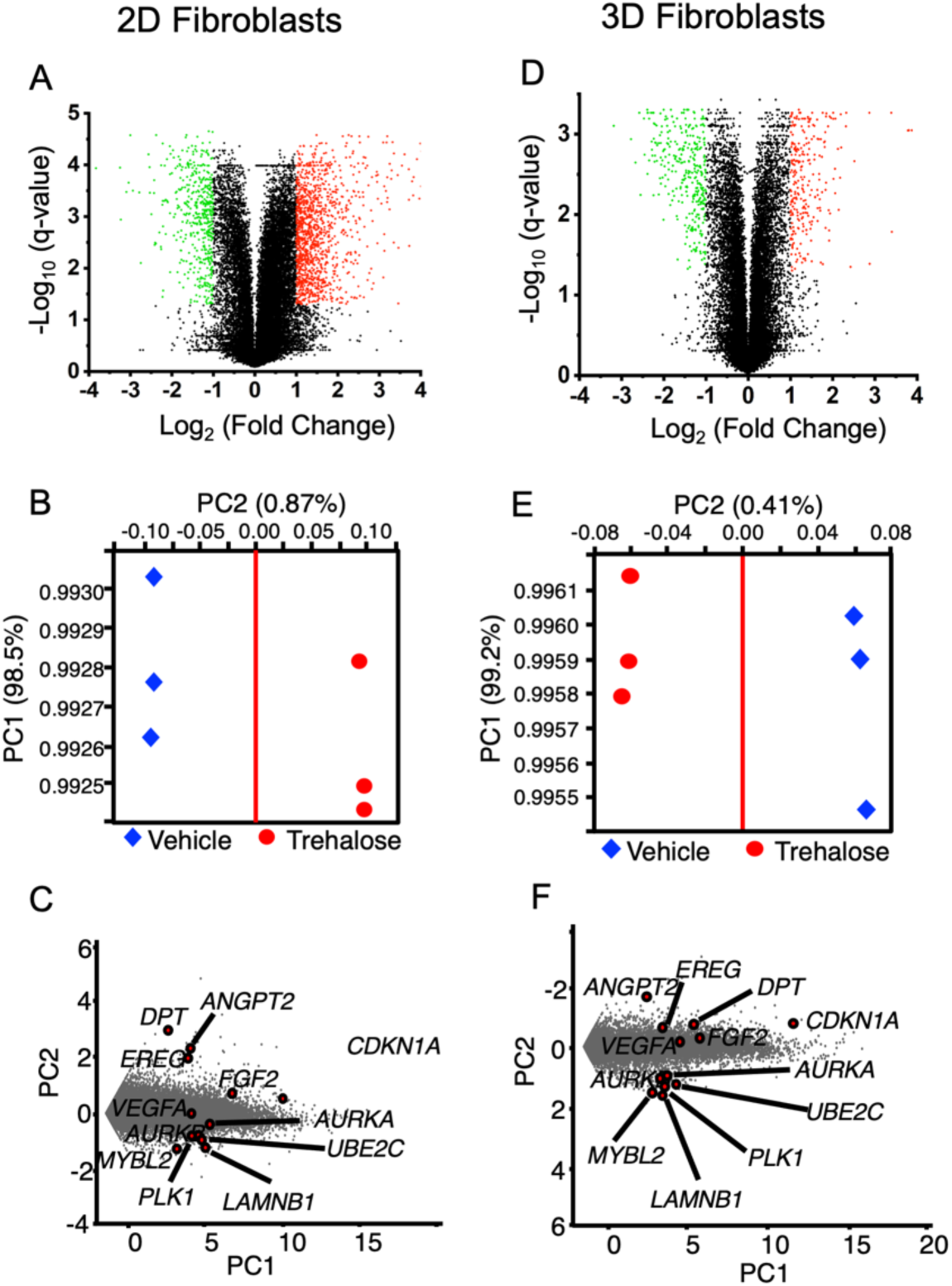
Gene expressions in highly concentrated trehalose-treated 2D and 3D fibroblasts by whole transcriptome analysis with RNA-seq. Volcano plots show that in the presence of trehalose (100 mg/ml) in 2D fibroblasts (**A**) and 3D fibroblasts (**D**), gene expressions were significantly modulated. Red or green rounds indicate genes increased more than 2-fold or decreased by less than the half, respectively, with less than 0.05 of q-values. PCA showing the separation between PC1 and PC2 in 2D fibroblasts (**B**) and 3D fibroblasts (**E**). The factor loadings of PC1 and PC2 of the genes calculated by PCA were plotted. The plotted upper and lower genes were detected in 2D fibroblasts (**C**) or 3D fibroblasts (**F**).

**Figure 3-figure supplement 1:**
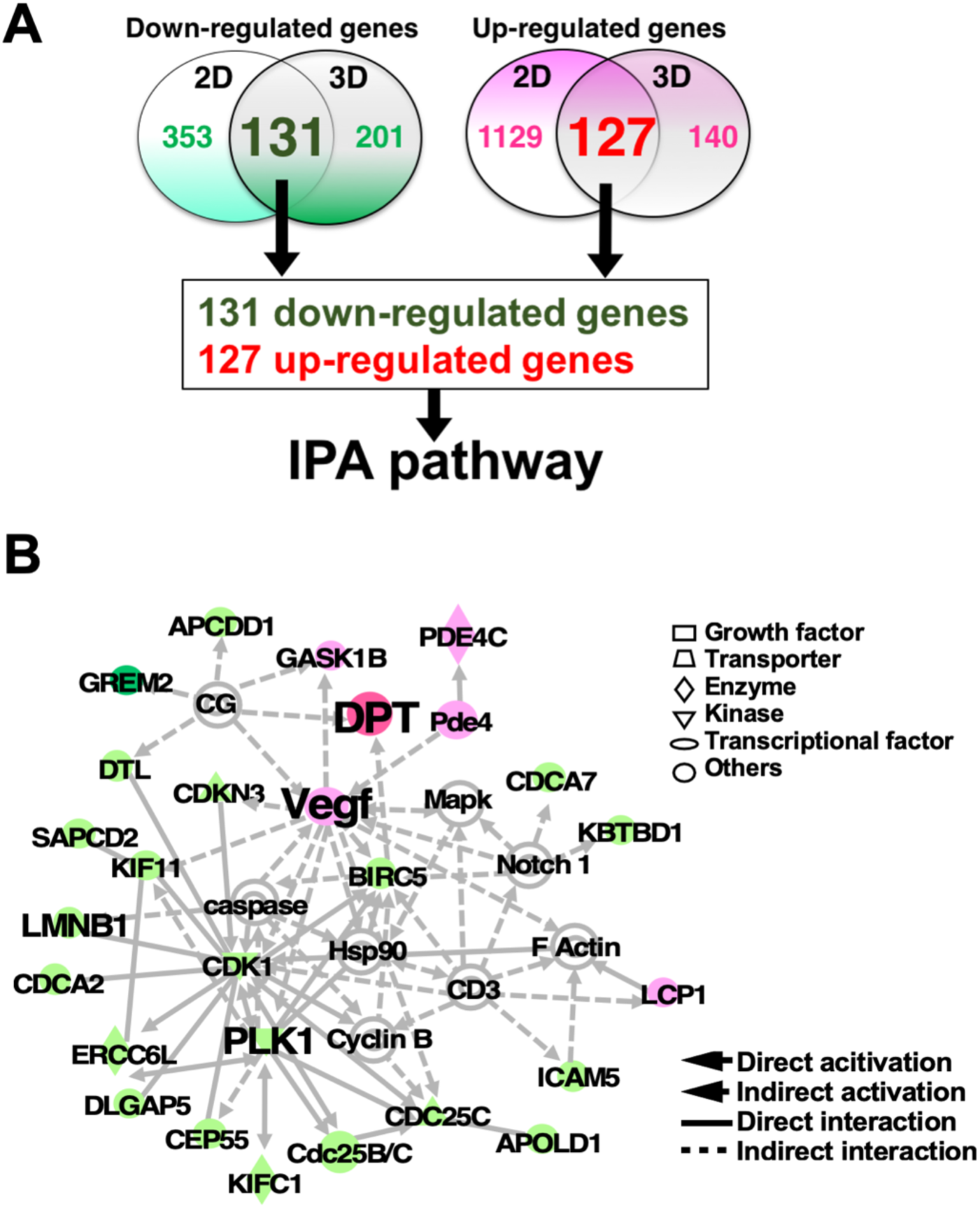
Network analysis by IPA pathway using genes modulated by trehalose. (**A**) The Venn diagram demonstrates the numbers of the downregulated genes and the upregulated-genes analyzed in Fig. 2. The 131 downregulated-genes and the 127 upregulated-genes were used for an IPA. (**B**) A network detected by IPA demonstrated activation of DPT and Vegf, accompanied by inhibition of PLK1, LMNB1, and CDK1. The red and green shapes demonstrate upregulated and downregulated genes by trehalose, respectively.

**Figure 4-figure supplement 1:**
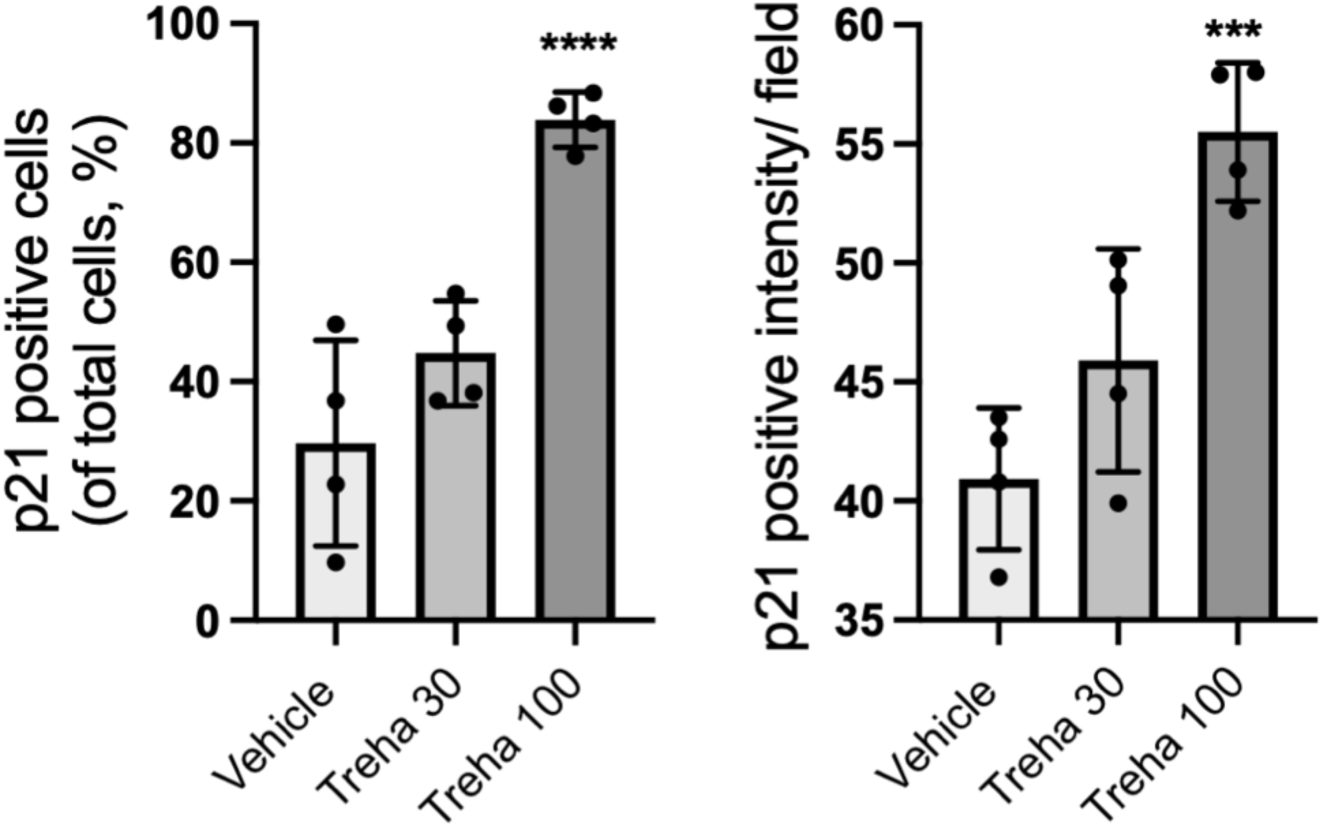
Trehalose modulates the expression of p21. Human dermal fibroblasts were treated with trehalose (30 and 100 mg/ml) or a vehicle control (PBS) for 24 hours. The cells were stained with antibody for p21 and DAPI for nuclei, and both were observed using a fluorescence microscope. In each group, we observed the relative number and intensity of fibroblasts stained by p21 antibody. Data are means ± SD for four wells, and are representative of three experiments with similar results. ***: *P* < 0.001, ****: *P* < 0.0001 versus the control (vehicle-treated) fibroblasts by one-way ANOVA).

**Figure 5-figure supplement 1:**
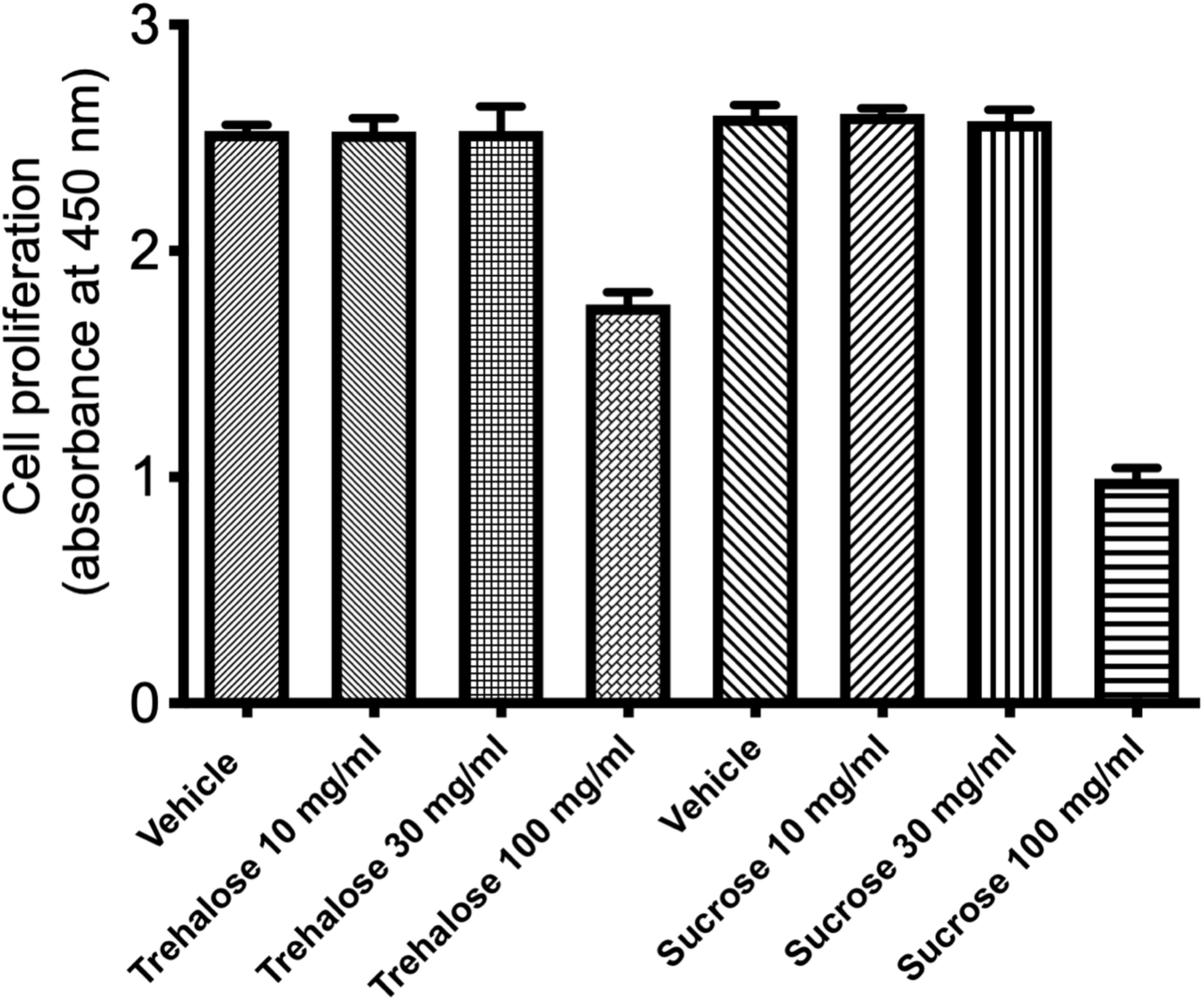
Proliferation assay with human dermal fibroblasts treated with trehalose. Proliferation assay with human dermal fibroblasts treated with trehalose (10, 30, and 100 mg/ml) or vehicle for 24 h.

**Figure 5-figure supplement 2:**
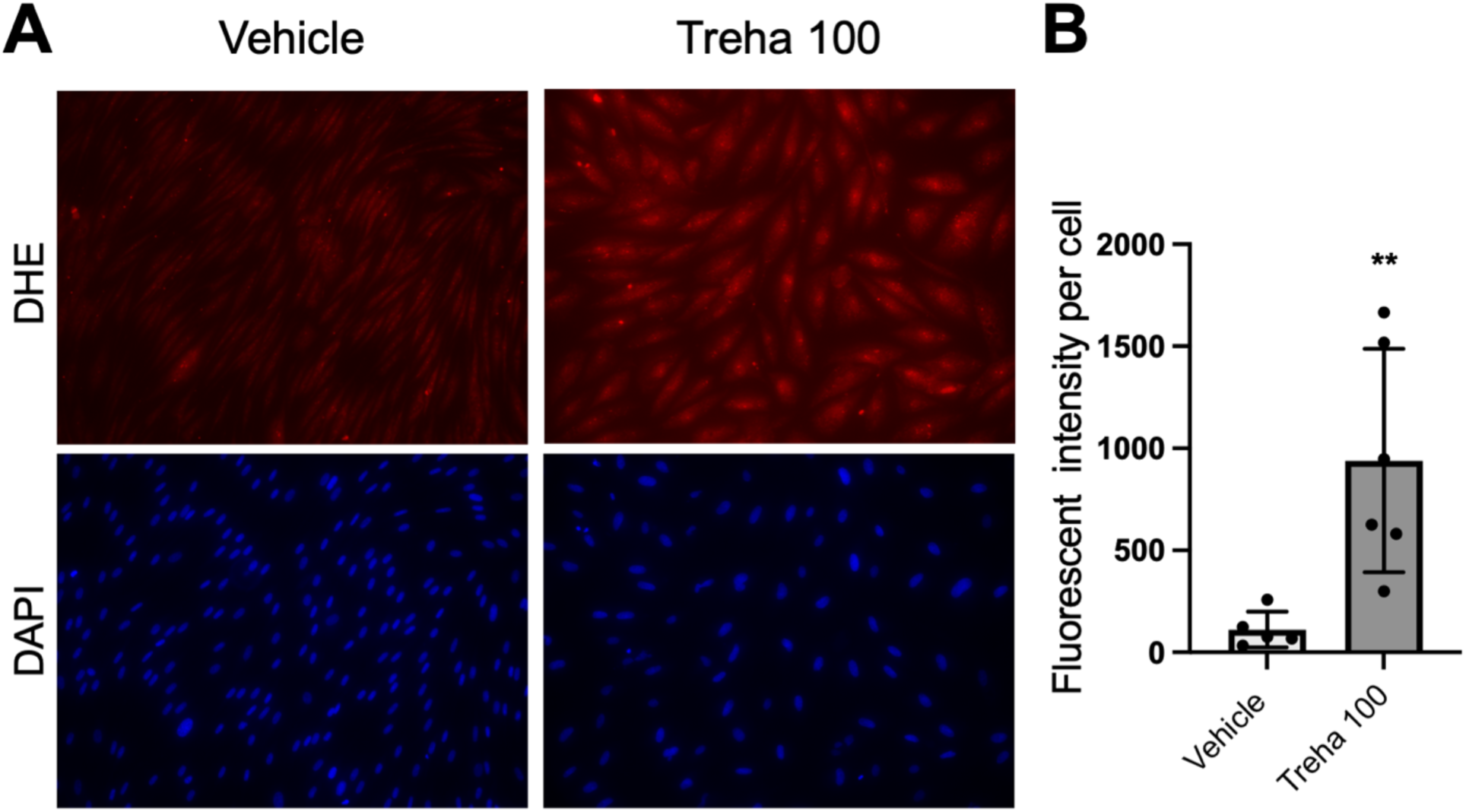
Fluorescent images of trehalose-treated fibroblasts stained with DHE. (**A**) Fibroblasts were treated with vehicle or trehalose (100 mg/ml) for 48 hours and stained with DAPI for nuclei. Representative fluorescent images of DHE-treated cells were obtained by fluorescence microscope. (**B**) In each group, we observed the intensity per cell stained with DHE. Data are means ± SD for four wells and are representative of two experiments with similar results. **: *P* < 0.001 versus the control (vehicle-treated) fibroblasts by Student *t*-test.

**Video 1.** The morphological alterations of the human dermal fibroblasts after vehicle treatment. Phase contrast microscopy imaging of the fibroblasts cultured with vehicle.

**Video 2.** The morphological alterations of the human dermal fibroblasts after the trehalose treatment (30 mg/ml). Phase contrast microscopy imaging of the fibroblasts cultured with trehalose.

**Video 3.** The morphological alterations of the human dermal fibroblasts after the trehalose treatment (100 mg/ml). Phase contrast microscopy imaging of the fibroblasts cultured with trehalose.

**Video 4.** The morphological alterations of the human dermal fibroblasts after the sucrose treatment (100 mg/ml). Phase contrast microscopy imaging of the fibroblasts cultured with sucrose.

